# Protein composition of axonal dopamine release sites in the striatum

**DOI:** 10.1101/2022.08.31.505994

**Authors:** Lauren Kershberg, Aditi Banerjee, Pascal S. Kaeser

## Abstract

Mechanisms of neuromodulatory transmission in the brain remain ambiguous. Dopamine is a prototypical neuromodulator, and it was recently found that its secretion relies on active zone-like release site assemblies. Here, we use in vivo biotin-identification (iBioID) proximity proteomics in mouse striatum to isolate dopamine release site proteins enriched over the general dopamine axonal protein content. Using three bait proteins, we identified 527 proteins that fall into several synaptic protein classes, including active zone, Ca^2+^ regulatory and synaptic vesicle proteins. We also detected many proteins not previously associated with synaptic exocytosis. Knockout of the presynaptic organizer protein RIM profoundly disrupted dopamine release site composition assessed by iBioID, while Synaptotagmin-1 knockout did not. α-Synuclein, a protein linked to Parkinson’s disease, was enriched at release sites, and this enrichment was lost in both tested mutants. We conclude that RIM organizes scaffolded dopamine release sites and we define the protein composition of these sites.

## Introduction

The molecular mechanisms of synaptic transmission have been studied extensively. In contrast, mechanistic understanding of neuromodulatory signaling lags behind. In the brain, dopamine is a critical neuromodulator for movement and reward, and dopamine operates through G-protein-coupled receptor (GPCR) signaling like most modulatory transmission (Berke, 2018; Liu et al., 2021a; Surmeier et al., 2014). The predominant source of dopaminergic innervation in the vertebrate brain stems from midbrain dopamine neurons. The dopamine neuron somata are located in the substantia nigra pars compacta (SNc) and the ventral tegmental area (VTA) and send prominent axonal projections to the striatum. In the striatum, dopamine is thought to act as a volume transmitter because only a small percentage of dopamine varicosities are directly apposed to post-synaptic specializations (Descarries et al., 1996; Wildenberg et al., 2021) and because dopamine receptors are found mostly extrasynaptically (Liu et al., 2021a; Rice et al., 2011; Sesack et al., 1994; Uchigashima et al., 2016; Yung et al., 1995). Hence, dopamine likely signals by diffusing through the striatal extracellular space before initiating responses in target cells. While much is understood about the mechanisms and molecules involved in the synaptic release of classical neurotransmitters, the machinery involved in release of volume transmitters such as dopamine is less well known.

Spatial and temporal precision of vesicular exocytosis at fast synapses is established by the active zone, a conserved molecular machine that docks synaptic vesicles at the presynaptic plasma membrane close to voltage-gated Ca^2+^ channels, primes the vesicles for release, and aligns these processes with postsynaptic receptor clusters (Biederer et al., 2017; Südhof, 2012). Several lines of evidence point to the existence of active zone-like protein complexes for the control of striatal dopamine release. First, dopamine release is rapid, has a high release probability, and occurs at small hotspots (Banerjee et al., 2020, 2022; Beyene et al., 2019; Liu et al., 2018; Marcott et al., 2014; Patriarchi et al., 2018; Silm et al., 2019; Wang et al., 2014; Zych and Ford, 2022), indicating the need for vesicle tethering close to Ca^2+^ channels before the stimulus arrives. Second, at least some dopamine receptors respond rapidly to local, high concentrations of dopamine (Beckstead et al., 2004; Condon et al., 2021; Courtney and Ford, 2014; Gantz et al., 2013; Marcott et al., 2014), which necessitates synchrony of vesicular release. Third, dopamine neuron activity often correlates with behavior on fast time scales (Bova et al., 2020; Chaudhury et al., 2013; Hamilos et al., 2021; Hollerman and Schultz, 1998; Howe and Dombeck, 2016; Jin and Costa, 2010; Schultz et al., 1997; da Silva et al., 2018), suggesting the need for precise signaling mechanisms. Taken together these properties indicate the need for protein machinery for rapid and efficient release of dopamine.

Indeed, our recent work has found that proteins for the control of spatial and temporal precision of synaptic vesicle exocytosis are important for evoked dopamine release. These proteins include the active zone scaffolds RIM and Liprin-α, the priming protein Munc13, and the fast Ca^2+^ sensor Synaptotagmin-1 (Syt-1) (Banerjee et al., 2020, 2022; Liu et al., 2018; Robinson et al., 2019). Conversely, the active zone scaffolds RIM-BP and ELKS are dispensable for axonal dopamine release, even though at least ELKS is present at these sites (Banerjee et al., 2022; Liu et al., 2018). Furthermore, striatal dopamine release only partially depends on voltage-gated Ca^2+^ channels of the CaV2 family, which are required for stimulated synaptic release (Brimblecombe et al., 2015; Held et al., 2020; Liu et al., 2022; Luebke et al., 1993; Takahashi and Momiyama, 1993). Thus, there appear to be both similarities and differences in the secretory machines at classical synapses and in dopamine varicosities. However, only a handful of proteins important for secretion are known to mediate dopamine release, and many pieces of the underlying secretory machine remain unidentified.

We here characterized the composition of dopamine release sites using an unbiased proteomic approach. We used in vivo biotin-identification (iBioID) of proteins, a method which allows for purifying proteins in close proximity to a marker protein (Kim et al., 2014; Roux et al., 2012; Uezu et al., 2016), to define the molecular composition of striatal dopamine release sites. An inclusive list of the dopamine release site proteome, generated with three different bait proteins, contains a total 527 proteins, with 190 proteins enriched in multiple bait conditions and 41 proteins enriched with all three baits over the general dopamine axonal protein content. Dopamine release sites contain many proteins with previously known functions at classical synapses including active zone proteins, Ca^2+^ regulatory proteins and synaptic vesicle proteins, as well as proteins that presently do not have defined roles in axonal neurotransmitter secretion. We also tested whether structural scaffolding and dopamine release are important for the composition of these secretory sites. Conditional, dopamine neuron-specific knockout of the presynaptic organizer protein RIM strongly disrupted the release site proteome, while knockout of the fast Ca^2+^ sensor to abolish synchronous dopamine release did not have strong effects on release site composition. We conclude that dopamine release sites are organized structures which are controlled by the scaffolding protein RIM, and we define the protein content of these dopamine release sites.

## Results

### Proximity proteomics to assess dopamine release site composition in the mouse striatum

With the overall goal to generate a comprehensive proteome of release sites in striatal dopamine axons, we adapted a proximity proteomic approach (Uezu et al., 2016). We fused the biotinylase BirA to several proteins associated with dopamine release sites and expressed these BirA bait proteins using Cre-dependent AAVs specifically in dopamine neurons of DAT^IRES-Cre^ mice to locally biotinylate proteins in vivo, followed by affinity purification of the biotinylated proteins and mass spectrometry for protein identification (Fig. 1A). The striatal dopamine axons are particularly well suited for this approach because they are elaborately branched in the striatum and can be easily separated from their midbrain somata and dendrites during tissue dissection, avoiding confounds arising from the co-purification of somatodendritic proteins.

**Figure 1.**
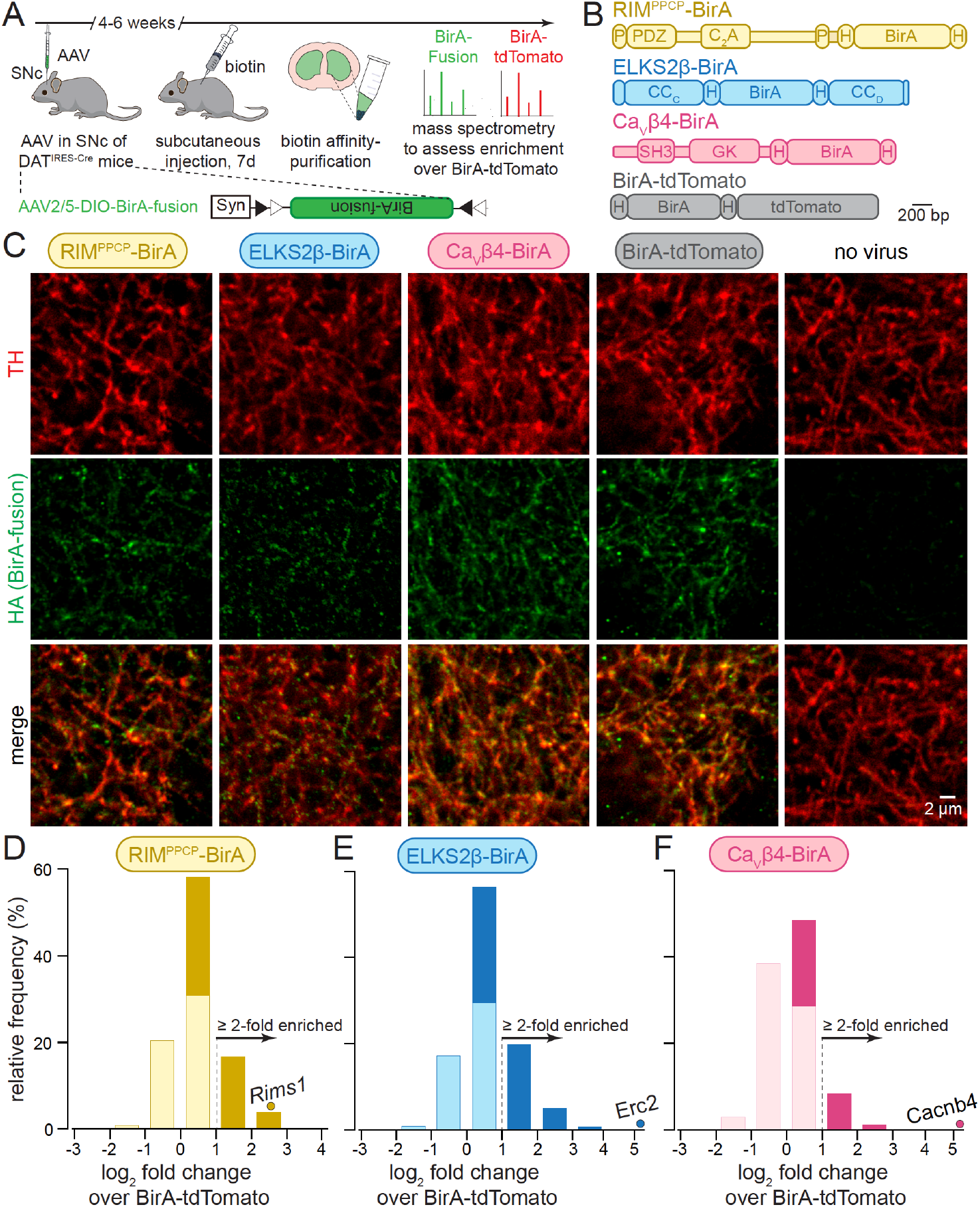
iBioID for release site proteins in dopamine axons of the mouse striatum. **(A)** Schematic of the iBioID experiment for purification of release site proteins from striatal dopamine axons using Cre-dependent AAVs (AAV2/5-DIO-BirA-fusion) and in vivo biotinylation followed by affinity purification and analyses by mass spectrometry. **(B)** Overview of BirA fusion proteins expressed via Cre-dependent AAVs in midbrain dopamine neurons of DAT^IRES-Cre^ mice, P: proline rich region, H: hemagglutinin (HA) tag. Each mouse expressed one of the three BirA baits (RIM^PPCP^-BirA, ELKS2β-BirA and CaVβ4-BirA) or BirA-tdTomato (to generate a dopamine axonal control proteome). **(C)** Representative confocal images of brain slices of DAT^IRES-Cre^ mice expressing the BirA fusion proteins shown in B, slices were stained with anti-TH antibodies and anti-HA antibodies to label dopamine axons and BirA fusion proteins, respectively. Images are from an experiment in which all constructs were imaged in the same session and with the same settings, and adjusted identically for display, except for ELKS2β-BirA. Images for ELKS2β-BirA were acquired in a separate experiment and with its own control, and images were adjusted slightly differently to match overall appearance, and adjustments were cross-checked against the control run with ELKS2β-BirA. Representative example of three to five images per mouse and condition, each experiment was repeated in ≥ 3 mice. **(D-F)** Protein enrichment in BirA bait conditions over BirA-tdTomato. Log_2_ fold change values are plotted as frequency histograms. Values at or below 0 represent proteins that are lower than in the BirA-tdTomato condition (light colors), values > 0 represent proteins that are higher than in the BirA-tdTomato condition (saturated colors), and values ≥ 1 represent hits (≥ 2-fold enriched); D, 1306 total proteins identified, 659 proteins with log_2_ fold change > 0, and 269 proteins with log_2_ fold change ≥ 1 (hits); E, 1496/805/382; F, 1168/354/114. The gene encoding the protein that was used as a bait is shown as a dot in each panel; D, 4 independent repeats (12 striata each); E, 4 (12/12/12/10 striata); F, 4 (12/12/12/10 striata). For pilot iBioID experiments and assessment of self-biotinylation, see Fig. S1.

We first generated a series of Cre-dependent AAVs to express BirA bait proteins of ELKS or RIM, active zone proteins associated with dopamine release sites (Banerjee et al., 2022; Liu et al., 2018), or of the β4 subunit of voltage-gated Ca^2+^ channels (CaVβ4), a component of CaV Ca^2+^ channel complexes that are present at release sites (Brimblecombe et al., 2015; Dolphin, 2003; Held et al., 2020; Tan et al., 2022a). Considering the packaging limit of AAVs and to avoid disrupting protein localization or function by BirA tagging, the following constructs were used (Fig. 1B): ELKS2β, a short, endogenous version of ELKS with the tag inserted between the CCC and CCD domains at a locus that does not disrupt ELKS function (ELKS2β-BirA) (Held et al., 2016; Kaeser et al., 2009; Liu et al., 2014); the central region of RIM that is important for its active zone localization (Tan et al., 2022a; Wu et al., 2019) and includes the PDZ and C2A domains and the linker sequences between them, flanked on both sides by two proline rich motifs, with the tag in a position that tolerates insertion without disrupting RIM function (RIM^PPCP^-BirA) (Kaeser et al., 2011; Tan et al., 2022a; Tang et al., 2016); and the CaVβ4 subunit with the tag inserted at the C-terminus, a locus that does not disrupt CaVβ4 localization (Held et al., 2020; Tan et al., 2022a). In all experiments, we used BirA fused to tdTomato that localizes throughout the axon as a control. All constructs also contained hemagglutinin (HA)-tags flanking BirA for identification with HA antibodies.

To assess the axonal presence of these BirA fusion proteins, we injected these Cre-dependent AAVs into the midbrain of DAT^IRES-Cre^ mice. After six weeks, we perfused the mice and stained striatal brain slices with antibodies against HA (to label the fusion proteins) and tyrosine hydroxylase (TH, to label dopamine axons) (Fig. 1C). The fusion proteins were detected within TH-labelled axons. RIM^PPCP^-BirA and ELKS2β-BirA were present in small punctate structures consistent with release site localization in dopamine axons (Liu et al., 2018). CaVβ4 was somewhat more wide-spread, similar to Ca^2+^ entry in these axons (Pereira et al., 2016), and BirA-tdTomato was broadly overlapping with TH, likely reflecting a soluble axonal localization. Obtaining high-quality super-resolved images from these stainings, similar to what we presented before for endogenous dopamine axonal proteins (Banerjee et al., 2020, 2022; Liu et al., 2018), was not possible, likely because of the specific antibody combinations that were required here. In vivo biotinylation was next confirmed for the BirA fusion proteins with or without biotin injections (for seven days) followed by pilot biotin-affinity purifications and Western blotting of the purified fractions (Fig. S1 and Materials and Methods). This established that biotinylation and protein purification is efficient, and that background biotinylation in the absence of biotin is negligible.

To determine dopamine release site composition, we performed iBioID using these three BirA bait proteins and BirA-tdTomato expressed in DAT^IRES-Cre^ mice (Fig. 1A). After four to six weeks of expression, biotin injections were done on seven consecutive days, and 10 to 12 striata per condition and repeat were dissected and homogenized. Biotinylated proteins were then isolated using affinity purification. The eluates were depleted for the two endogenously biotinylated proteins pyruvate carboxylase (PC) and propionyl-CoA carboxylase (PCCA) before assessment of protein content with mass spectrometry. In total, four independent biological repeats per BirA fusion protein were performed (that is, four times 10-12 striata per condition, conditions: RIM^PPCP^-BirA, ELKS2β-BirA, CaVβ4-BirA, BirA-tdTomato, also see Materials and Methods). The four repeats were run as two large experiments for in vivo biotinylation, biotin affinity purification and mass spectrometry; two repeats were run in parallel in each experiment, and the two experiments were run approximately one year apart from one another (the second experiment also contained the analyses of conditional knockout mice described below). Peptides from 1969 proteins were identified in at least one of the BirA conditions. 300 (15%) of those proteins are mitochondrial proteins, which is common in iBioID experiments, and these proteins were removed from further analyses unless noted otherwise, as has been done before (Loh et al., 2016; Takano et al., 2020; Uezu et al., 2016). To identify proteins enriched in the immediate vicinity of the bait proteins (ELKS2β-BirA, RIM^PPCP^-BirA, CaVβ4-BirA), the number of peptides identified for each protein and bait was normalized to the number of peptides found for the same protein in the BirA-tdTomato condition, effectively generating a ratio of enrichment over soluble dopamine axonal protein content as “fold change”. For proteins that were only detected with the bait, but not with BirA-tdTomato, we used a value of 0.5 peptides for BirA-tdTomato for calculating fold-change. The log_2_ of the fold change was then produced, with values > 0 representing proteins detected at levels higher than in the BirA-tdTomato condition (Figs. 1D-1F, saturated colors), while proteins ≤ 0 (light colors) were below. “Hits” were proteins enriched ≥ 2-fold and have log_2_ fold change values of ≥ 1 (Figs. 1D-1F); these proteins are considered enriched and included in further analyses (Fig. 2).

**Figure 2.**
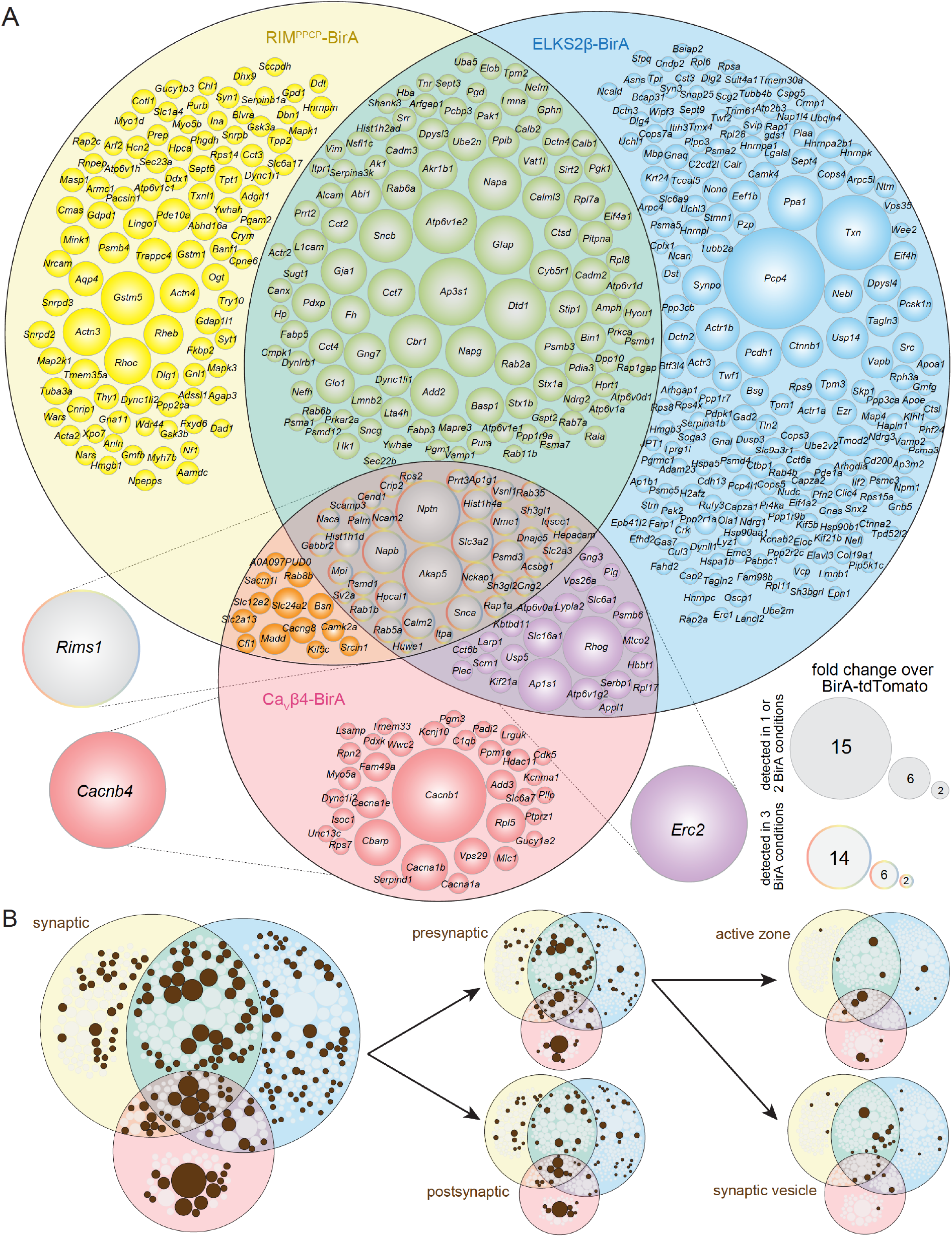
The protein composition of release sites in dopamine axons. **(A)** Venn diagram listing genes that encode proteins enriched at least 2-fold over BirA-tdTomato with RIM^PPCP^-BirA (yellow), ELKS2β-BirA (blue) or CaVβ4-BirA (red) iBioID baits. Proteins used as BirA baits are shown outside the Venn diagram in the color of the corresponding part of the diagram. Enrichment is shown as the average of four independent repeats run in two mass spectrometry sessions (repeats 1 and 2 in the first session, repeats 3 and 4 in the second) and was analyzed as described in the methods. Circle size reflects average enrichment across repeats (key on the bottom right). For proteins enriched in multiple bait conditions, circle size corresponds to the bait condition with the largest enrichment. **(B)** Enriched proteins from (A) that have cellular compartment annotations in the SynGO database (Koopmans et al., 2019) are colored in brown and classified into increasingly specific SynGO sub-categories, sub-categorizations are not mutually exclusive. For assessment of Neuroplastin (*Nptn*) localization in striatal synaptosomes, see Fig. S2.

For RIM^PPCP^-BirA (Fig. 1D) and ELKS2β-BirA (Fig. 1E), 50% and 54% of the detected proteins were higher than with BirA-tdTomato, and 20% and 25% were above the two-fold enrichment threshold, respectively. Only 30% of all proteins detected in the CaVβ4-BirA condition were higher compared to BirA-tdTomato, and 9% were enriched ≥ 2-fold. These observations align with the morphological data (Fig. 1C), where CaVβ4-BirA was more widely expressed than RIM^PPCP^ and ELKS2β-BirA. They are also consistent with functional analyses that revealed that Ca^2+^ entry in dopamine axons is widespread (Pereira et al., 2016), while active zone proteins (Banerjee et al., 2022; Liu et al., 2018) and vesicle fusion events (Pereira et al., 2016) are only detected in 20 to 30% of the varicosities. A key feature in each condition is that the protein that was used as a BirA bait was highly enriched (individual circles in Figs. 1D-1F). This further establishes the specificity of the approach as self-biotinylation of the BirA fusion protein should be high compared to other proteins if the labeling radius is small, which has been estimated to be 10 to 50 nm (Kim et al., 2014; Roux et al., 2012).

### Assessment of the protein composition of release sites in dopamine axons

We next constructed Venn diagrams with proteins that were above the ≥ 2-fold enrichment threshold to assess the overlap of hits between the different bait conditions. In total, there were 527 proteins ≥ 2-fold enriched (Fig. 2A). The enrichment compared to BirA-tdTomato indicates that these proteins have a localization that is restricted compared to diffusible dopamine axonal proteins. Of the 527 proteins, 190 were enriched in more than one condition and 41 were enriched with all three baits, RIM^PPCP^-BirA, ELKS2β-BirA and CaVβ4-BirA.

Given that the BirA baits are proteins with roles in transmitter secretion and are expressed in dopamine axons, it is expected that proteins with known roles in neurotransmitter release are enriched in the iBioID dataset. We assessed the hits by evaluating their cellular compartment annotations in SynGO, an expert-curated database that assigns localization and function of synaptic genes (Koopmans et al., 2019). Of the 527 enriched proteins, 196 (37%) have one or multiple synaptic cellular compartment annotations in SynGO (Fig. 2B), reflecting that their synaptic localization has been established before. SynGO does not distinguish between synapse or neurotransmitter type, but instead broadly determines whether there is evidence to support the presence or function of a given protein at synapses in general (Koopmans et al., 2019).

To gain a deeper understanding of the types of synaptic proteins enriched in the dataset, the proteins were categorized based on whether they are known to be localized pre- or postsynaptically. Proteins with SynGO annotations may have multiple annotations and can be both pre- and postsynaptic, but do not require a more specific annotation if evidence supporting a more specific subsynaptic localization is insufficient. 103 proteins in the dataset (19.5% of the entire dataset, 52.5% of the SynGO synaptic proteins) have a presynaptic annotation with many having additional specific assignments to the active zone or synaptic vesicles, while 88 (16.7% of the entire dataset, 44.5% of the SynGO synaptic proteins) have postsynaptic annotations (Fig. 2B). 43 proteins (7.8% of the entire dataset, 21.9% of the SynGO synaptic proteins) are annotated both pre- and postsynaptically.

If the iBioID approach used here enriched proteins specifically at sites of dopamine release, it would be expected that proteins previously shown to localize to dopamine release sites are enriched. Indeed, RIM (*Rims1*), ELKS (*Erc2*) and Bassoon (*Bsn*), proteins that can be used as markers for dopamine release sites (Banerjee et al., 2022; Liu et al., 2018), were all enriched across multiple conditions (Fig. 2A).

A key advantage of proteomic assessment of release site composition, like the one described here, is the identification of putative new candidates. Of the 527 enriched proteins, 331 (63%) do not have a SynGO annotation (Figure 2B), and 46 proteins are annotated broadly as “synaptic” without more exact specification. Additionally, for the characterization of the dopamine axon secretory machinery, proteins known to be at classical synaptic release sites that have not been implicated in dopamine release can also be identified. One example of the latter category is Neuroplastin (*Nptn*), a transmembrane protein with pre- and postsynaptic roles at classical synapses (Beesley et al., 2014; Boyken et al., 2013; Schmidt et al., 2017; Smalla et al., 2000). Neuroplastin is strongly enriched across conditions (Fig. 2A), but its presence in dopamine axons has not been described. To assess Neuroplastin localization in dopamine axons with a secondary approach, we prepared striatal synaptosomes as we described before (Banerjee et al., 2020, 2022; Liu et al., 2018) and stained them with antibodies against TH to mark dopamine synaptosomes, Bassoon to label release sites, and Neuroplastin. Neuroplastin staining intensity was significantly greater in TH-labeled synaptosomes that contained Bassoon than in those without Bassoon (Fig. S2). These results reveal that Neuroplastin is enriched in proximity to release sites of dopamine axons and validate iBioID for one of the main hits.

### Ablation of the active zone protein RIM, but not of the Ca^2+^ sensor Synaptotagmin-1, disrupts release site composition in dopamine axons

We next asked whether removing proteins important for dopamine release affects the composition of dopamine axon release sites. Previous work has established genetic strategies to abolish evoked dopamine release by removing the presynaptic scaffolding protein RIM (Banerjee et al., 2022; Liu et al., 2018; Robinson et al., 2019) or synchronous release by removing the fast Ca^2+^ sensor Syt-1 (Banerjee et al., 2020). We crossed mice with floxed alleles for RIM1 and RIM2 or for Syt-1 to DAT^IRES-Cre^ mice (Backman et al., 2006; Kaeser et al., 2011; Liu et al., 2018) to remove these proteins from dopamine neurons (Fig. 3A, RIM cKO^DA^ or Syt-1 cKO^DA^, respectively). We then performed iBioID analogous to the proteome described in Figs. 1 and 2 (called control from this point forward), with the modification that in each mutant we did two independent repeats instead of four (that is, two times 10 to 12 striata for each of the bait conditions in RIM cKO^DA^ and Syt-1 cKO^DA^ mice) due to the genetic complexity and volume of the experiment.

**Figure 3.**
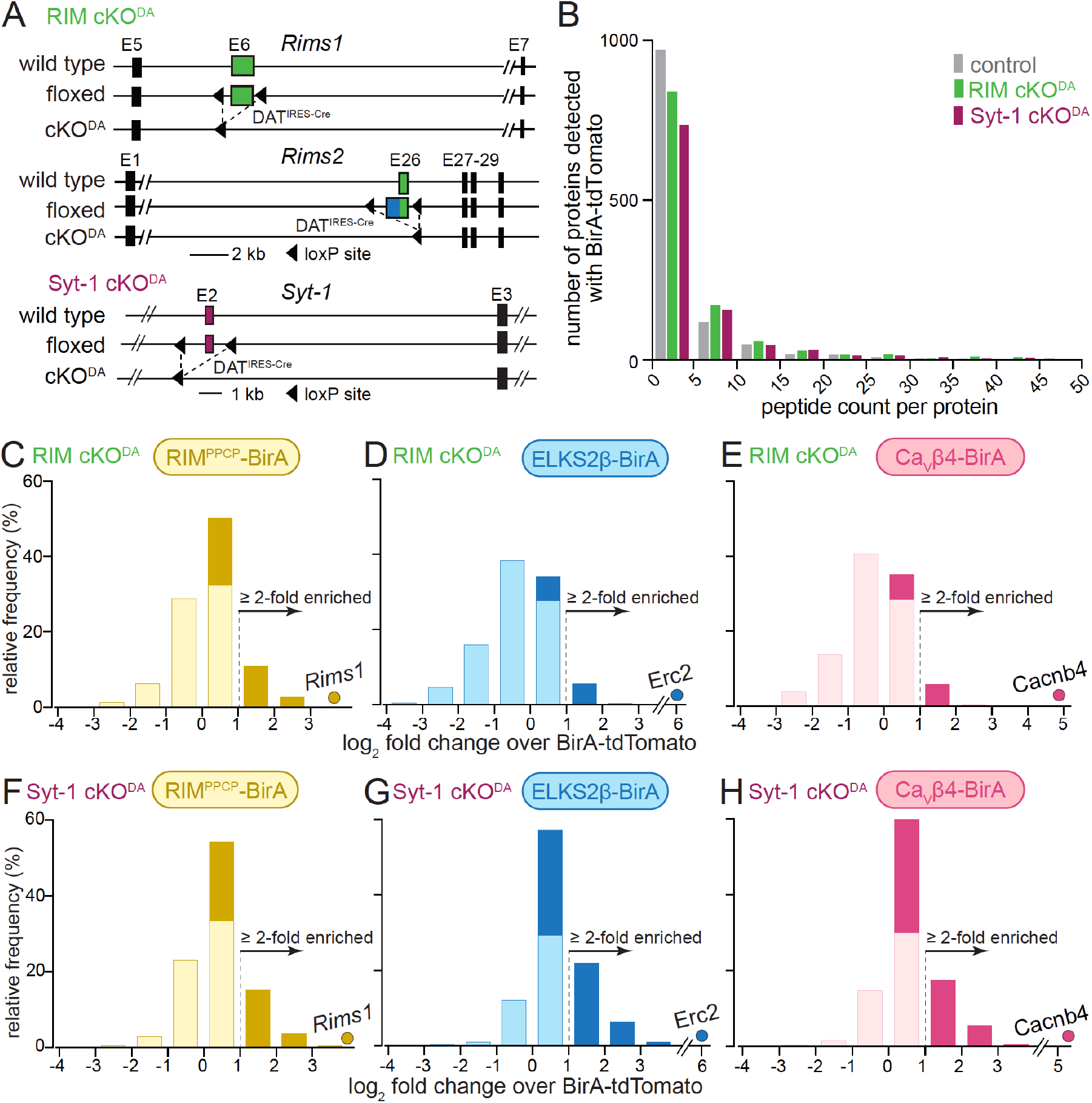
Enrichment of release site proteins after conditional ablation of RIM or of Synaptotagmin-1 in dopamine neurons. **(A)** Strategy for ablation of RIM (RIM cKO^DA^) or Synaptotagmin-1 (Syt-1 cKO^DA^) from dopamine neurons using conditional mouse genetics (Banerjee et al., 2020; Liu et al., 2018) **(B)** The average number of peptides in bins of five found for each protein with BirA-tdTomato for each genotype. The control is from data in Figs. 1 and 2 and is the average peptide count of four repeats, the proteomes from RIM cKO^DA^ and Syt-1 cKO^DA^ is an average peptide count of two repeats. The x-axis is cut at a peptide count of 50 covering > 99% of the detected proteins. Average number of detected proteins: control, 1195; RIM cKO^DA^, 1155; Syt-1 cKO^DA^, 1011. **(C-E)** Protein enrichment in BirA bait conditions over BirA-tdTomato in RIM cKO^DA^ mice. Log_2_ fold change values are plotted as frequency histograms. Values at or below 0 represent proteins that are lower than in the BirA-tdTomato condition (light colors), values > 0 represent proteins that are higher than in the BirA-tdTomato condition (saturated colors), and values ≥ 1 represent hits (≥ 2-fold enriched); C, 1089 total proteins identified, 334 proteins with log_2_ fold change > 0, and 149 proteins with log_2_ fold change ≥ 1 (hits); D, 1003/123/62; E, 1008/130/65. The gene encoding the protein that was used as a bait is shown as a dot in each panel, C, 2 independent repeats (10/12 striata); D, 2 (12/10); E, 2 (10/12). **(F-H)** Same as C-E, but for Syt-1 cKO^DA^ mice; F, 1017 total proteins identified, 424 proteins with log_2_ fold change > 0, and 199 proteins with log_2_ fold change ≥ 1 (hits); G, 1103/619/327; H, 1016/544/246; F, 2 independent repeats (10/12 striata); G, 2 (12/12); H, 2 (12/12). For Venn diagrams of release site protein enrichment, see Fig. S3 for RIM cKO^DA^, and Fig. S4 for Syt-1 cKO^DA^.

We first compared the BirA-tdTomato condition in control, RIM cKO^DA^ and Syt-1 cKO^DA^ and found that both the number of identified proteins and the peptide counts were similar across experiments (Fig. 3B). Hence, removal of RIM or Syt-1 from dopamine axons did not cause strong disruptions in the overall dopamine axon protein content in the striatum of these mutant mice, and similar amounts of axonal material were present, consistent with previous work that found TH axon density to be similar to control mice in these mutants (Banerjee et al., 2020; Liu et al., 2018).

We next assessed fold change over tdTomato for each of the BirA baits and each mutant. In RIM cKO^DA^ mice, 31% (RIM^PPCP^-BirA), 12% (ELKS2β-BirA) and 13% (CaVβ4-BirA) of the identified proteins were higher than with BirA-tdTomato, and 14% (RIM^PPCP^-BirA), 6% (ELKS2β-BirA) and 4% (CaVβ4-BirA) reached the ≥ 2-fold enrichment threshold to be considered hits (Fig. 3C-3E). These percentages are overall lower than in the control proteome (Figs. 1D-1F). The protein enrichment over BirA-tdTomato in Syt-1 cKO^DA^ mice was more similar to the control proteome (Figs. 3F-3H, RIM^PPCP^-BirA: 42% higher than with BirA-tdTomato and 20% ≥ 2-fold enriched; ELKS2β-BirA: 56% and 30%, CaVβ4-BirA 53% and 24%). We conclude that in RIM cKO^DA^ mice, protein enrichment at release sites in dopamine axons is disrupted compared to control or Syt-1 cKO^DA^ mice, suggesting that RIM removal disrupts mechanisms important for release site scaffolding. Abolishing synchronous dopamine release alone by Syt-1 cKO^DA^ has no strong effects on the protein composition of release sites in dopamine axons.

We next assessed hits in more detail by generating and analyzing Venn diagrams for each mutant. The RIM cKO^DA^ dataset contained a total of 198 hits (Fig. S3), compared to the 527 hits in the control proteome (Fig. 2). RIM1 was absent from all conditions except for RIM^PPCP^-BirA (which likely reflects self-biotinylation), confirming the efficacy of the conditional gene knockout strategy. It is noteworthy that 49% of all hits (98 out of 198) in the RIM cKO^DA^ dataset came from the RIM^PPCP^-BirA bait only (Fig. S3, yellow circle), indicating that re-expression of RIM^PPCP^, the central region of RIM containing its scaffolding domains (Fig. 1B), may restore some scaffolding deficits caused by RIM cKO^DA^. In Syt-1 cKO^DA^ mice, 450 hits were detected (Fig. S4), similar to the 527 hits in the control dataset, and 104 (23%) of the hits were at least two-fold enriched for all three BirA baits, including RIM (Fig. S4).

To characterize which proteins are depleted from the release site proteome in the RIM cKO^DA^ mice, we first used SynGO to categorize all identified proteins in each dataset into synaptic proteins (Fig. 4A, grey bars), presynaptic proteins (Fig. 4B) and active zone proteins (Fig. 4C). We then plotted the log_2_ of the fold change of the average of all annotated proteins in each category. In the control dataset (grey bars), synaptic proteins were enriched as expected (Fig. 4A), and the extent of enrichment had a tendency to increase as the SynGO annotation became more specific (to presynaptic, Fig. 4B, and to active zone, Fig. 4C). These effects not only disappeared in RIM cKO^DA^ mice, but instead reverted (Figs. 4A-4C, green bars), and for ELKS2β-BirA and CaVβ4-BirA, proteins in all three SynGO categories were depleted (log_2_ fold change values < 0). The depletion is absent in the RIM^PPCP^-BirA condition in RIM cKO^DA^ mice, supporting that release site scaffolding is partially restored. In contrast to the RIM cKO^DA^ dataset, the proteins found in the Syt-1 cKO^DA^ dataset show an enrichment that is overall relatively similar to the control dataset with all conditions having log_2_ fold change values > 0 (Figs. 4A-4C, maroon bars), supporting that abolishing synchronous dopamine release does not disrupt overall release site composition.

**Figure 4.**
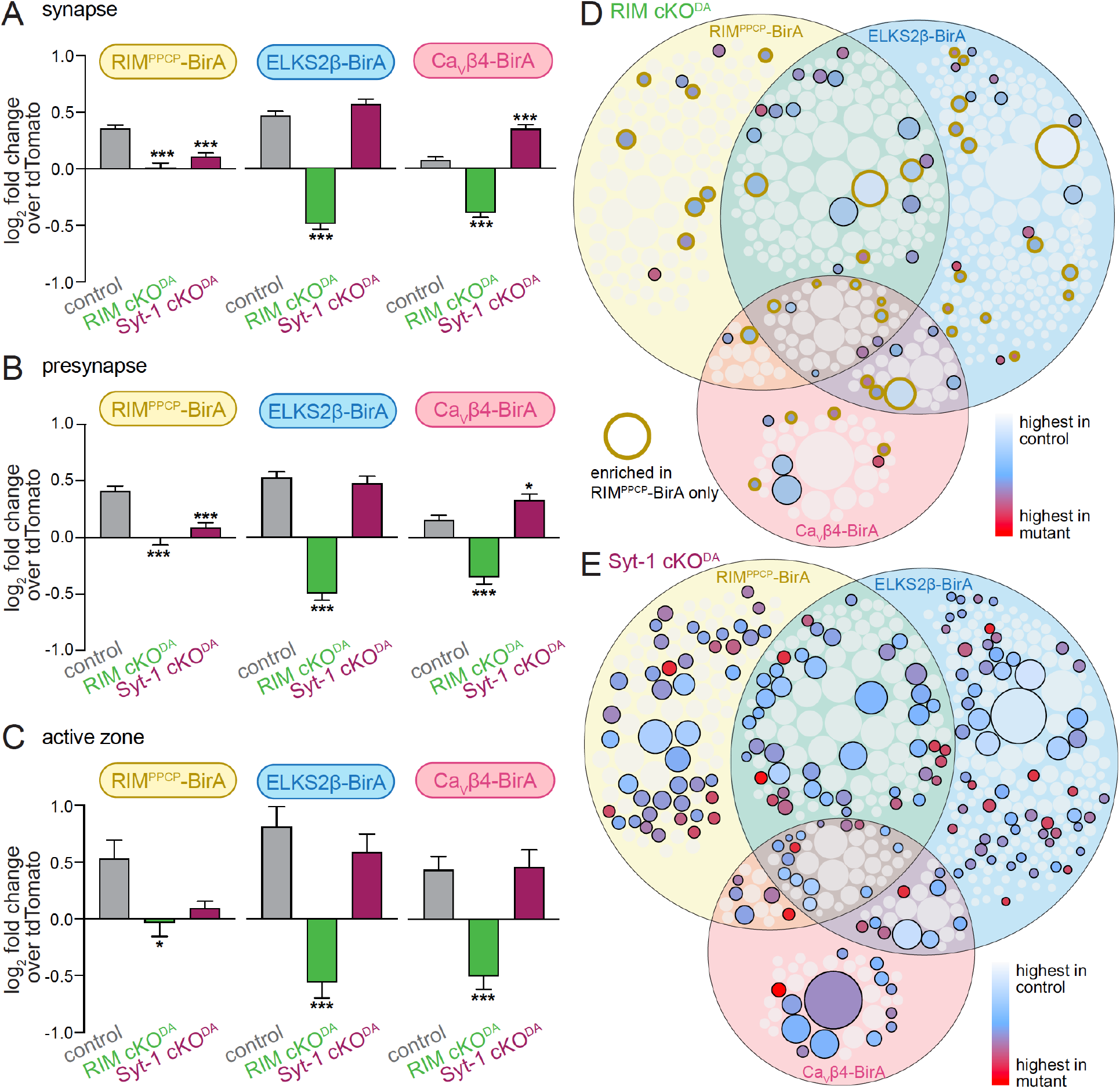
RIM cKO^DA^ strongly disrupts the protein composition of release sites in dopamine axons. **(A)** The average log_2_ fold change of identified proteins over the same proteins in the BirA-tdTomato conditions, all proteins with a synaptic localization annotation in SynGO for each bait and genotype are included. Positive values represent release site enrichment and negative values represent depletion relative to axonal protein content assessed with BirA-tdTomato. Total number of proteins detected with a SynGO annotation: control, 540; RIM cKO^DA^, 412; Syt-1 cKO^DA^, 438. **(B)** Same as A, but for proteins with a “presynapse” SynGO annotation. Total number of proteins: control, 257; RIM cKO^DA^, 208; Syt-1 cKO^DA^, 213. **(C)** Same as A, but only proteins an “active zone” SynGO annotation. Total number of proteins: control, 42; RIM cKO^DA^, 37; Syt-1 cKO^DA^, 36. **(D)** Proteins enriched in the RIM cKO^DA^ dataset mapped onto the control Venn diagram shown in Fig. 2A. Proteins present in the control proteome but not in the RIM cKO^DA^ dataset are shown in grey. Enriched proteins are colored based on relative enrichment (key on the right). Proteins in the RIM cKO^DA^ dataset that are only enriched with the RIM^PPCP^-BirA bait are outlined in light brown. **(E)** Same as D, but for the Syt-1 cKO^DA^ dataset and without outlining a specific condition. Data in A-C are shown as mean ± SEM, and significance is presented as * p < 0.05, ** p < 0.01, and *** p < 0.001. Two-way ANOVA was used in A-C (A: genotype ***, bait ***, interaction ***; B: genotype ***, bait **, interaction ***; C: genotype ***, bait *, interaction not significant), and Bonferroni post-hoc tests (p values indicated in figure) were used to compare each genotype to control for each BirA bait.

We next mapped the enriched proteins detected in each mutant onto the Venn diagram of the control proteome. Only 15% of proteins in the control dataset were also enriched in the RIM cKO^DA^ dataset (Fig. 4D), and 49% of those proteins stem from the RIM^PPCP^ condition (Fig. 4D, circles with yellow outline). In contrast, 37% of the proteins in the control dataset were also enriched in the Syt-1 cKO^DA^ dataset (Fig. 4E). There might not be full overlap between the control and Syt-1 cKO^DA^ datasets because the approach is not saturating and does not detect all release site proteins, because the mutant datasets were derived from a smaller number of mice, or because of variability that is expected for these types of proteomic analyses.

Finally, to assess the organization of the proteins and their interactions in the various release site proteomes, we used the STRING database (Snel et al., 2000; Szklarczyk et al., 2019), which combines both theoretical predictions and empirical data to generate maps of protein-protein functional and physical interactions. We selected the proteins that were enriched and assigned as synaptic proteins by SynGO (Fig. 2B, Fig. S3B, Fig. S4B, “synaptic”) and analyzed each dataset (Fig. 5A-5C). In the control dataset, the proteins formed an integrated network with functional nodes classified as active zone proteins, synaptic vesicle proteins, Ca^2+^ regulatory proteins and synaptic ribosomes (Fig. 5A), and the same nodes were detected in the Syt-1 cKO^DA^ dataset (Fig. 5C). In contrast, these networks were disrupted in the RIM cKO^DA^ dataset (Fig. 5B), even when hits from the RIM^PPCP^-BirA bait (lighter colors) are included. Only the synaptic ribosome node remained, indicating that while release site disruption is strong in RIM cKO^DA^ mice, other protein complexes near the bait proteins remain intact.

**Figure 5.**
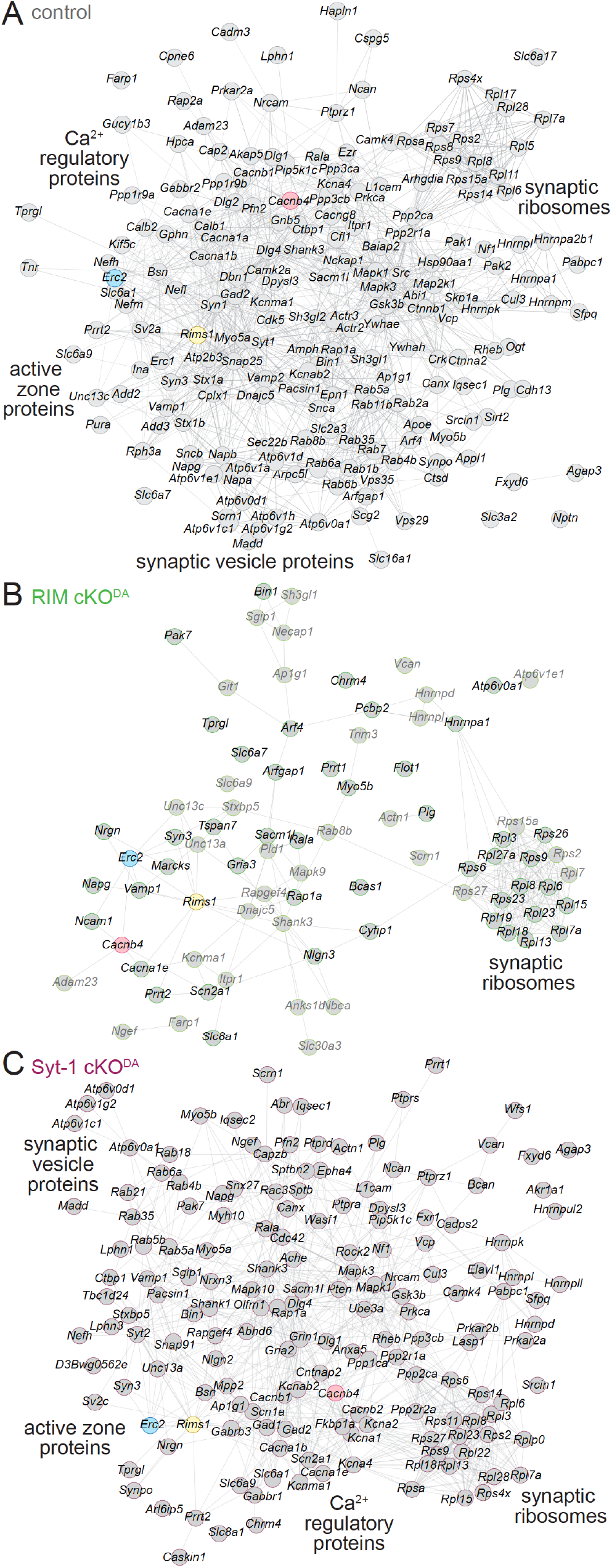
STRING diagrams illustrate functional categories of release site proteins and their disruption in RIM cKO^DA^ mice. **(A)** STRING diagram of the enriched proteins that have a synaptic SynGO annotation in the control dataset. Physical or functional interactions determined by either empirical data or predictive modeling are illustrated as lines between proteins (Snel et al., 2000; Szklarczyk et al., 2019). The BirA bait proteins are color-coded. Broad functional categories identified by STRING analyses are labeled close to the corresponding clusters. **(B)** As in A, but for the RIM cKO^DA^ dataset. The proteins that are enriched only in the RIM^PPCP^-BirA bait are shown in lighter grey. **(C)** As in A, but for the Syt-1 cKO^DA^ dataset.

### Disrupted recruitment of α-synuclein to release sites in dopamine axons after impairing dopamine release

Loss of dopamine neurons underlies the motor symptoms characteristic of Parkinson’s disease (Poewe et al., 2017), and recent human genetic studies indicate that mutations in *RIMS* genes increase the risk of Parkinson’s disease and are key drivers of disease progression (Liu et al., 2021b; Nalls et al., 2019). Monogenic forms of Parkinson’s disease exist and genetic associations with variable penetrance have been described (Blauwendraat et al., 2020; Day and Mullin, 2021; Marras et al., 2016). Our control dataset contained five of these genes: *Snca, Park7, Dnajc6, Synaptojanin-1*, and *Vps35* (Fig. 6A). Three were enriched in the control dataset with at least one of the BirA baits, and *Snca*, which encodes the synaptic vesicle associated protein α-synuclein, was above ≥ 2-fold enrichment threshold in each condition. In both RIM cKO^DA^ and Syt-1 cKO^DA^ mice, α-synuclein was substantially reduced and fell below the enrichment threshold (Fig. 6B). These data suggest that synchronous dopamine release or another activity that is shared between RIM and Syt-1 in dopamine axons is necessary for α-synuclein recruitment to release sites in these axons.

**Figure 6.**
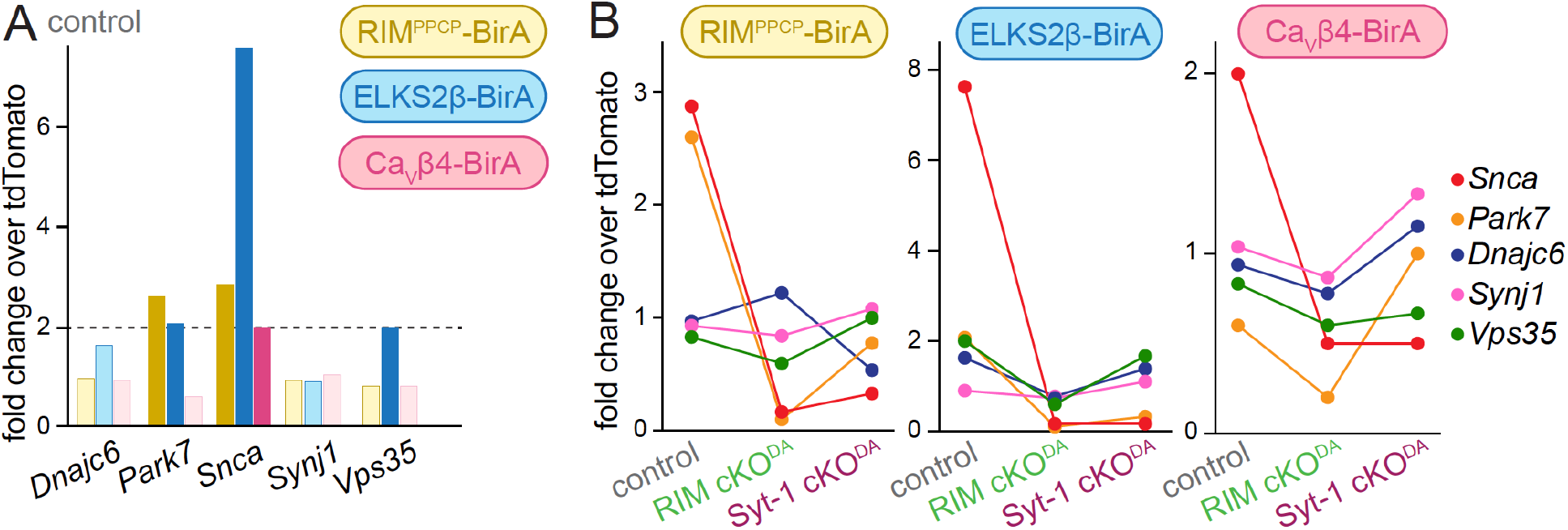
Genes associated with Parkinson’s disease in the dopamine release site dataset. **(A)** Proteins representing five Parkinson’s disease genes (Blauwendraat et al., 2020; Day and Mullin, 2021; Marras et al., 2016) were present in the control dataset, and their enrichment is plotted for each condition. Hits above the two-fold enrichment threshold are shown in saturated colors, and those below the threshold in light colors; control, 4 independent repeats per condition (as described in Fig. 1). **(B)** Enrichment of the five proteins shown for each bait and in control, RIM cKO^DA^ and Syt-1 cKO^DA^ conditions; control, 4 independent repeats (as described in Fig. 1), RIM cKO^DA^, 2 (as described in Fig. 3); Syt-1 cKO^DA^, 2 (as described in Fig. 3).

## Discussion

We used iBioID to assess the composition of release sites in dopamine axons. Three different bait proteins associated with dopamine release sites were fused to BirA and specifically expressed in striatal dopamine axons. After in vivo biotinylation and affinity purification, we identified 527 proteins enriched in the immediate vicinity of the secretory proteins over dopamine axonal proteins, and 190 of those proteins were enriched in multiple bait conditions. The list of enriched proteins was composed of known secretory machinery including classical active zone proteins, synaptic vesicle proteins, Ca^2+^ regulatory proteins, and additional proteins not previously associated with vesicular exocytosis. Conditional knockout of the presynaptic organizer protein RIM from dopamine neurons caused a strong disruption of release site composition assessed with iBioID, with loss of most of the enriched proteins compared to the control proteome. In contrast, removal of the vesicular Ca^2+^ sensor for synchronous dopamine release, Syt-1, did not cause major changes using the same approach. We conclude that striatal dopamine axons contain active zone-like sites with secretory machinery for rapid vesicular exocytosis, present a proteomic assessment of the molecular composition of this site, and show that its integrity strongly depends on the scaffolding mechanisms of RIM.

### Proximity proteomics establish a central scaffolding role for RIM in dopamine axons

In previous studies, we have assessed roles of individual active zone proteins in striatal dopamine release (Banerjee et al., 2022; Liu et al., 2018). Several active zone proteins were found to be present in release site-like structures in dopamine axons, including Bassoon, RIM, Munc13, and ELKS. Using conditional knockout, we established that RIM and Munc13 are essential for evoked axonal dopamine release, while ELKS and RIM-BP are dispensable. RIM removal from dopamine neurons also induced deficits in the clustering of Bassoon in striatal dopamine axons. Based on these findings, we proposed the model of an active zone-like release site in dopamine axons for action potential-evoked dopamine release (Banerjee et al., 2022; Liu and Kaeser, 2019; Liu et al., 2018, 2021a).

It is not possible to generate a molecular map of the dopamine secretory pathway with candidate approaches, and it has remained uncertain whether RIM is important for active zone scaffold assembly, or whether RIM is only essential for dopamine exocytosis but is not a key tethering factor. We adapted a proximity proteomic approach to generate a list of components that may be localized to and function at secretory hotspots in dopamine axons. This list contains several classes of proteins including active zone proteins, synaptic vesicle proteins, Ca^2+^ regulatory proteins, and many additional proteins. A key feature across datasets is the strong enrichment of RIM, perhaps indicating a central role in dopamine active zone assembly. In strong support of this model, RIM was essential for dopamine release site composition assessed by iBioID. RIM ablation from dopamine axons resulted in a strong reduction in the number of proteins identified as release site components. This loss of protein material is illustrated more strongly in the ELKS2β-BirA and CaVβ4-BirA conditions than with RIM^PPCP^-BirA, suggesting that re-expression of the RIM^PPCP^-BirA may partially restore release site composition.

Altogether, our data indicate that removal of RIM causes disruption of dopamine release site structures. It validates the iBioID approach because many of the proteins in the control proteome were lost upon RIM knockout, suggesting that enrichment is specific. We cannot conclusively rule out that some of the effect of loss of release site proteins in RIM cKO^DA^ mice arises from mis-localized BirA bait proteins in dopamine axons. Several lines of evidence, however, suggest that aberrant localization is unlikely to fully explain the disruption of the release site proteome: (1) Twenty-two proteins are enriched with all three bait proteins in the RIM cKO^DA^ mice, suggesting overlapping BirA labeling radii with BirA baits localized near one another in the RIM cKO^DA^ mice (Fig. S3A). (2) The identified ribosome cluster is unaffected in the RIM cKO^DA^ dataset (Fig. 5), suggesting that some of the proteome is intact and colocalizes with all three baits. (3) Finally, previous studies suggest that overall dopamine axon structure in RIM cKO^DA^ is not detectably impaired (Liu et al., 2018) and the axonal proteome identified by BirA-tdTomato was similar between RIM cKO^DA^ and wild type axons (Fig. 3). Overall, these points make it unlikely that mislocalization of the three BirA baits explains disruption of the release site proteome in RIM cKO^DA^ axons. Nevertheless, if all three baits were mislocalized, this would further support the overall conclusion that knockout of RIM in dopamine axons disrupts release site structure. Moreover, and from a functional perspective, it is impossible to assess whether the BirA baits are localized to release sites in RIM cKO^DA^ mice because RIM knockout abolishes evoked dopamine release, and hence there are no functional release sites left.

*RIMS1* has been identified as a risk factor and *RIMS2* as a disease progression gene for Parkinson’s disease in human genetic studies (Liu et al., 2021b; Nalls et al., 2019). Five proteins from the monogenic forms of Parkinson’s disease (Blauwendraat et al., 2020; Day and Mullin, 2021; Marras et al., 2016) were detected in dopamine axons using BirA-tdTomato, and three of those proteins exceeded the enrichment threshold for at least one bait protein (Fig. 6). α-Synuclein was enriched with all BirA baits and ablation of RIM or Syt-1 disrupted α-synuclein enrichment. α-Synuclein is associated with synaptic vesicles and regulates synaptic vesicle clustering, SNARE complex formation, and endocytosis (Burré et al., 2010; Diao et al., 2013; Murphy et al., 2000). RIM or Syt-1 knockout causes a loss of docked vesicles at synapses (Chang et al., 2018; Kaeser et al., 2011), possibly explaining the loss of α-synuclein enrichment at release sites of dopamine varicosities in RIM cKO^DA^ and Syt-1 cKO^DA^ mice. The loss of release site enrichment of α-synuclein in dopamine axons when RIM is removed highlights that classical release site proteins may be associated with brain disease through roles in neuromodulatory transmission. Overall, our findings might support a model in which RIM is associated with Parkinson’s disease through dopamine release site recruitment of α-synuclein, possibly via protein interactions or through vesicle docking.

### An active zone-like site for dopamine release

Two recent studies have assessed general aspects of the composition of dopamine axons or of dopamine varicosities using different methodologies. One study used mass spectrometry to analyze dopaminergic synaptosomes obtained through fluorescent sorting (Paget-Blanc et al., 2022) and identified 57 proteins in dopaminergic synaptosomes, which contain release sites, vesicles, mitochondria, cytoskeletal elements and many other components. A second study used a proximity proteomic approach with APEX2 to identify axonal proteins (Hobson et al., 2022), similar to the BirA-tdTomato condition used for normalization in our experiments. These approaches identified proteins overlapping with our current (Figs. 1-6) and previous (Banerjee et al., 2022; Liu et al., 2018) work, but neither of the two studies was designed to enrich for release site proteins over other axonal proteins. Notably, there is good overlap between axonal proteins in our study and the previous study that used biotinylation, with 65% of the proteins identified in our axonal control proteome (BirA-tdTomato condition) also present in dopamine axonal proteome published in (Hobson et al., 2022).

We defined the inclusion criterion for release site association as ≥ 2-fold enrichment over axonal proteins identified via BirA-tdTomato, a cutoff similar to other iBioID studies (Takano et al., 2020; Uezu et al., 2016). We detected multiple classes of proteins beyond classical active zone scaffolds (Figs. 2, 5). Synaptic vesicle proteins, for example, were highly abundant, indicating that synaptic vesicles are enriched at release sites compared to the axonal cytosol, likely by being tethered in a fusion-competent state. This is consistent with an overall high vesicular release probability for evoked dopamine secretion and with the strong dependence of dopamine release on RIM and Munc13 (Banerjee et al., 2022; Liu et al., 2018), proteins that dock and prime vesicles at conventional synapses (Kaeser and Regehr, 2017; Südhof, 2012).

Which active zone-associated proteins are identified in our release site proteomes? RIM (*Rims1*), Bassoon (*Bsn)*, and ELKS1 (*Erc1*), ELKS2 (*Erc2*), and P/Q-type (CaV2.1, *Cacna1a*) and N-type Ca^2+^ channels (CaV2.2, *Cacna1b*) were all robustly detected, consistent with previous studies that assessed roles and/or localization of these proteins in dopamine neurons (Banerjee et al., 2022; Brimblecombe et al., 2015; Daniel et al., 2009; Ducrot et al., 2021; Liu et al., 2018, 2022; Uchigashima et al., 2016). Notably absent from the proteins in the enriched release site dataset are the active zone proteins Liprin-α and RIM-BP (Emperador-Melero and Kaeser, 2020; Schoch and Gundelfinger, 2006; Südhof, 2012). Removal of Liprin-α2 and -α3 in dopamine neurons has less severe effects on dopamine release compared to RIM or Munc13 knockout, while removal of RIM-BP leaves dopamine release unimpaired (Banerjee et al., 2022; Liu et al., 2018). Thus, the abundance of active zone proteins in our dataset correlates overall with the strength of their functional roles in dopamine release or their established presence in dopamine axons. One exception to this overall correlation is Munc13. We recently found that evoked striatal dopamine release is strongly impaired after Munc13 ablation (Banerjee et al., 2022). In the proteomic datasets, Munc13 was present, but only enriched in some cases (Figs. 2, 5, S3 and S4); it did not consistently appear as a strongly enriched protein. At synapses, some Munc13 may be more broadly distributed than just at active zones (Tan et al., 2022b, 2022a), but this does not explain its non-enrichment here because in the BirA-tdTomato dataset, Munc13 was not strongly detected either. Hence, Munc13 may either be overall sparse despite its requirement for evoked dopamine release, or may be difficult to detect, for example because it is poorly biotinylated or not easily purified in biotin pulldowns. Overall, the low presence of Munc13 and the partial overlap between release site proteomes of control and Syt-1 cKO^DA^ mice illustrate that our work does not reveal a saturated, full proteome, but some proteins may be below detection threshold, as is common for proteomic experiments.

Dopamine neurons corelease the fast neurotransmitters glutamate and GABA (Hnasko et al., 2010; Stuber et al., 2010; Tritsch et al., 2012). While GABA is loaded into vesicles via VMAT2 and hence co-released from the same vesicular compartment (Melani and Tritsch, 2022; Tritsch et al., 2012), glutamate co-release may or may not be occurring at the same release sites (Hnasko et al., 2010; Silm et al., 2019; Zhang et al., 2015). Some of the secretory machinery detected in the iBioID proteomes may mediate release of fast transmitters from dopamine neurons, and it is currently not possible to determine whether they might depend on dedicated release sites with specific molecular machinery. Overall, however, our results fit well with previous candidate approaches on dopamine release site proteins (Banerjee et al., 2022; Daniel et al., 2009; Liu et al., 2018), suggesting that the proteome we describe represents dopamine release sites to a significant extent.

### Postsynaptic and unknown proteins in the release site proteome from dopamine axons

There were 88 proteins with postsynaptic SynGO annotations in the release site proteome, and three possibilities may account for this. (1) The SynGO database may not have enough empirical evidence to designate any given protein as presynaptic. Some proteins with only postsynaptic annotations may be genuinely present in the dopamine axon release machinery. (2) Dopamine axons receive input from cholinergic interneurons (Cachope et al., 2012; Threlfell et al., 2012; Zhou et al., 2001), and activation of nicotinic receptors triggers dopamine axon action potential firing (Liu et al., 2022). The structure and organization of this cholinergic input is commonly considered non-synaptic (Chang, 1988; Jones et al., 2001), but recent evidence suggests that functionally, synaptic-like transmission exits (Kramer et al., 2022). Hence, dopamine varicosities may in principle contain postsynaptic scaffolds to tether neurotransmitter receptors, although nicotinic acetylcholine receptors were not enriched in the control dataset. (3) Biotinoyl-5’-AMP, generated by the BirA enzyme, has an estimated labeling radius of up to 50 nm (Kim et al., 2014; Roux et al., 2012). Striatal dopamine axons make sparse synaptic connections with medium spiny neurons that contain Gephyrin, a GABAergic postsynaptic scaffold (Uchigashima et al., 2016; Wildenberg et al., 2021). Gephyrin (*Gphn)* was indeed enriched with the ELKS2β-BirA and RIM^PPCP^-BirA baits (Fig. 2). Biotinylation across membranes has been observed before, for example for intramitochondrial proteins and with BirA-Gephyrin baits in which synaptic vesicle proteins were also enriched (Uezu et al., 2016). It is noteworthy that in our experiments, dopamine receptors were not enriched, consistent with dopamine receptor localization outside of the postsynaptic scaffolds in target neurons (Uchigashima et al., 2016).

The enriched dataset also contains 63% proteins that are not annotated in SynGO. These proteins may embody real hits that are not included in SynGO because there is not yet enough data to define them as presynaptic proteins. Some of these hits may be specific to dopamine release sites. Our work opens the door for future research to test whether any given protein is localized to and functions at dopamine release sites. It is possible that some hits arise because the BirA bait proteins are not exclusively at release sites. For example, it is clear that release site machinery including the baits need to be transported along the axon, and indeed several kinesins (*Kif21a, Kif21b, Kif5a, Kif5b)* were enriched. Hence, proteins of transport packets containing active zone and other proteins may be enriched over the BirA-tdTomato axonal proteome. The synaptic ribosomes (Fig. 5) that were unequivocally detected across conditions may also reflect some non-active zone-localized bait proteins. Finally, it is possible that some proteins are simply mis-identified as experimental noise.

## Conclusions

Taken together, the presented data and previous studies support a model of active zone-like dopamine release sites (Banerjee et al., 2022; Ducrot et al., 2021; Hobson et al., 2022; Liu and Kaeser, 2019; Liu et al., 2018, 2021a; Pereira et al., 2016). The significant loss of release site proteins when RIM is removed from dopamine neurons appears different from classical synapses, where removing RIM alone does not severely disrupt release site structure, but redundant scaffolding mechanisms have to be simultaneously disrupted (Acuna et al., 2016; Emperador-Melero and Kaeser, 2020; de Jong et al., 2018; Kaeser et al., 2011; Kushibiki et al., 2019; Oh et al., 2021; Tan et al., 2022b, 2022a; Wang et al., 2016). This supports the conclusion that dopamine release sites may have less redundancy in their scaffolding mechanisms and rely more heavily on RIM (Banerjee et al., 2022; Liu et al., 2018), and this central importance of RIM might explain why mutations in it are associated with Parkinson’s disease in human genetic studies (Liu et al., 2021b; Nalls et al., 2019).

## Acknowledgements

We thank J. Wang and C. Qiao for technical support, current and former members of the Kaeser laboratory and D. Brann for insightful discussions and feedback, M. Feany and H. Nyitrai for comments on the manuscript, R. Tomaino and the Taplin Mass Spectrometry Facility at Harvard Medical School for help with sample analyses, and S. Soderling and A. Uezu for sharing the BirA plasmid and for advice on in vivo biotinylation and biotin affinity purification early in the project. This work was supported by the NIH (R01NS103484, PSK; F31NS105159, LK), the Lefler foundation (to PSK), an HMS Dean’s Innovation grant (PSK), a Quan fellowship (LK), and an Alice Joseph Brooks fellowship (AB). We acknowledge the Neurobiology Imaging Facility (supported by a P30 Core Center Grant P30NS072030).

## Author contributions

Conceptualization, LK and PSK; Methodology, LK and AB; Formal Analysis, LK and PSK; Investigation, LK; Resources, LK and AB; Writing – Original Draft, LK and PSK; Writing – Review & Editing, LK, AB and PSK; Supervision, PSK; Funding Acquisition, PSK.

## Declaration of interests

The authors declare no competing interests. LK is currently an employee of Mass General Brigham (Boston, MA, USA) and was previously employed by the Prescient Healthcare Group (Jersey City, NJ, USA).

## Materials and Methods

### Key Resources Table

| Key Resources Table |  |  |  |  |
| --- | --- | --- | --- | --- |
| Reagent type (species) or resource | Designation | Source or reference | Identifiers | Additional information |
| genetic reagent (*M. musculus*) | *B6.SJL-Slc6a3<sup>tm1.1(Cre)Bkmm</sup>/J* (DAT<sup>IRES-Cre</sup>) | (Backman et al., 2006) | RRID:IMSR_JAX:006660 |  |
| genetic reagent (*M. musculus*) | *Rims1<sup>tm3Sud</sup>/J* (RIM1αβ<sup>fl/fl</sup>) | (Kaesler et al., 2008) | RRID:IMSR_JAX:015832 |  |
| genetic reagent (*M. musculus*) | *Rims2<sup>tm1.1Sud</sup>/J* (RIM2αβγ<sup>fl/fl</sup>) | (Kaesler et al., 2011) | RRID:IMSR_JAX:015833 |  |
| genetic reagent (*M. musculus*) | *C57BL/6Ntac-Syt1<sup>tm1a(EUCOMM)Wtsi</sup>/WtsiCnrm* (Syt-1<sup>fl/fl</sup>) | (Zhou et al., 2015) | RRID:IMSR_EM:06829 | The identifier refers to the line before flip recombination |
| cell line (*Homo sapiens*) | HEK293T cells | ATCC | Cat#: CRL-3216<br>RRID:CVCL_0063 |  |
| recombinant DNA reagent | pAAV2/5-syn-DIO-RIM-PRM-PDZ-PxxP-C2A-BirA | This study | LK18005; lab plasmid code (LPC) p864 |  |
| recombinant DNA reagent | pAAV2/5-syn-DIO-ELKS2β-BirA | This study | LK17008; LPC p857 |  |
| recombinant DNA reagent | AAV2/5-syn-DIO-Cavβ4-BirA | This study | LK19004; LPC p868 |  |
| recombinant DNA reagent | AAV2/5-syn-DIO-BirA-tdTomato | This study | LK17011; LPC p860 |  |
| antibody | anti-HA (rabbit polyclonal) | Cell Signaling Technology | CAT# 5017,<br>RRID:AB_10693385,<br>lab antibody code (LAC) A40 | Immunofluorescence (IF) (1:500) |
| antibody | anti-HA (mouse monoclonal) | Biolegend | CAT# 901501<br>RRID:AB_2565006;<br>LAC A12 | IF (1:500)<br>Western blot (WB) 1:500 |
| antibody | anti-tyrosine hydroxylase (guinea pig polyclonal) | SySy | CAT# 213 104,<br>RRID:AB_2619897,<br>LAC A111 | IF (1:1000) |
| antibody | anti-Bassoon (mouse monoclonal) | Millipore | CAT# ADI-VAM-PS003-F,<br>RRID:AB_11181058,<br>LAC A85 | IF (1:500) |
| antibody | anti-neuroplastin (goat polyclonal) | R and D systems | CAT# AF5360<br>RRID:AB_2155920<br>LAC A253 | IF (1:500) |
| antibody | anti-red fluorescent protein (rabbit polyclonal) | Rockland | CAT#600-401-379<br>RRID:AB_2209751,<br>LAC A81 | WB (1:1000) |
| antibody | anti-β-actin (mouse monoclonal) | Sigma Aldrich | CAT# A1978<br>RRID:AB_476692<br>LAC A127 | WB (1:10000) |
| antibody | anti-pyruvate carboxylase (rabbit polyclonal) | Novus | CAT# NBP1-49536G<br>RRID:AB_11016707<br>LAC A252 |  |
| software, algorithm | Prism9 | Graphpad | RRID:SCR002798<br>https://www.graphpad.com/scientific-software/prism |  |
| software, algorithm | Cytoscape v 3.8.2 | Cytoscape | RRID:SCR003032<br>http://cytoscape.org/ |  |
| software, algorithm | SEQUEST PRO | Thermo Fisher | https://scicrunch.org/resolver/SCR_014594 |  |
| software, algorithm | MATLAB code for object analysis | (Liu, 2020) | https://github.com/kaeserlab/3DSIM_Analysis_CL |  |

### Mouse lines

Expression of AAVs and conditional deletion of active zone proteins in dopamine neurons was performed using DAT^IRES-Cre^ mice (Backman et al., 2006) (Jackson laboratories; RRID:IMSR_JAX: 006660, *B6.SJL-Slc6a3^tm1.1(Cre)Bkmm^*/J). iBioID to generate the control proteome was performed in mice heterozygote for DAT^IRES-Cre^. Conditional ablation of RIM in dopamine neurons (RIM cKO^DA^) was performed as previously described (Liu et al., 2018). Mice with floxed alleles for *Rims1* (to remove RIM1α and RIM1β, RRID:IMSR_JAX:015832, *Rims1^tm3Sud^*/J) (Kaeser et al., 2008) and *Rims2* (to remove RIM2α, RIM2β and RIM2γ, RRID:IMSR_JAX:015833, *Rims2^tm1.1Su^d*/J) (Kaeser et al., 2011) were crossed to DAT^IRES-Cre^ mice. Conditional ablation of Syt-1 in dopamine neurons (Syt-1 cKO^DA^) was performed as previously described (Banerjee et al., 2020). Mice with floxed alleles for *Syt-1* were previously produced from a gene targeting experiment at EMMA (RRID:IMSR_EM:06829, *C57BL/6Ntac-Syt1^tm1a(EUCOMM)Wtsi^*/WtsiCnrm) and were flp recombined and crossed to DAT^IRES-Cre^ mice as described before (Banerjee et al., 2020; Zhou et al., 2015). For generating cohorts of mice for iBioID, the floxed alleles were homozygote in both parents and one parent contained a heterozygote DAT^IRES-Cre^ allele. Male and female mice were used in all experiments irrespective of sex. All animal experiments were approved by the Harvard University Animal Care and Use Committee (protocol number IS00000049).

### Cell lines and AAV production

AAVs were used to express BirA fusion proteins in dopamine neurons: LK18005-AAV2/5-syn-DIO-RIM^PPCP^-BirA (also called LK18005-AAV2/5-syn-DIO-RIM-PRM-PDZ-PxxP-C2A-BirA, p864), LK17008-AAV2/5-syn-DIO-ELKS2β-BirA (p857), LK19004-AAV2/5-syn-DIO-CaVβ4-BirA (p868), and LK17011-AAV2/5-syn-DIO-BirA-tdTomato (p860). All AAVs were made in HEK293T cells (purchased mycoplasma free from ATCC, CRL-3216, RRID:CVCL_0063, immortalized human cell line of female origin) using Ca^2+^ phosphate transfection. All AAVs were of the serotype 2/5. Three days after transfection, HEK293T cells were collected, and stored in freezing buffer (150 mM NaCl, 20 mM Tris-Cl, 2 mM MgCl2, pH 8.0) at -20 °C until viral purification. For purification of AAVs, cells were lysed by three freeze-thaw cycles with dry ice/ethanol and a 37 °C incubator. After 1 h benzonase nuclease treatment at 37 °C, cells were loaded onto an iodixanol gradient (5 ml each, 15%, 25%, 40%, 60%) and ultra-centrifuged at 208,000 x g for 4 h. Viral particles were then purified from the 40% layer of the gradient. Quantitative reverse transcriptase PCR was used to determine viral titers, and viruses were used at concentrations ranging from 4.0 x 10^11^ to 9.6 x 10^12^ viral genome copies/ml.

### Stereotaxic surgery and biotin injections

Mice (at postnatal days 30 to 55) were anesthetized in a 5% isoflurane induction chamber and then mounted on a stereotaxic frame, stable anesthesia was maintained with 1.5 to 2% isoflurane for the length of the surgery with a nose cone. The scalp was cut open and a hole was drilled in the skull and 1 µl of AAV viral solution was injected in the substantia nigra pars compacta (right or bilaterally depending on the experiment, 0.6 mm anterior, +/-1.3 mm lateral of Lambda and 4.2 mm below the surface of the brain) using a microinjector pump (PHD ULTRA, Harvard Apparatus) at 100 nl/min. Mice were treated with post-surgical analgesia and were allowed to recover for at least 28 days prior to biotin injections (for iBioID) or transcardial perfusion (for morphological experiments). Biotin injections were started four to six weeks after stereotaxic AAV injection. Mice were subcutaneously injected for seven consecutive days with 500 µl of 5 mM biotin dissolved in phosphate-buffered saline (PBS).

### Immunostaining and confocal imaging of brain sections

At least four weeks after stereotaxic injection of BirA viruses, mice (58 to 100 days old) were deeply anesthetized with isoflurane. Transcardial perfusion was performed with 30 to 50 ml ice-cold PBS followed by 50 ml of 4% paraformaldehyde in PBS (4% PFA) at 4 °C. Brains were then dissected out and incubated in 4% PFA for overnight at 4 °C. Fixed brains were sliced on a vibratome (Leica, VT1000s) at 100 μm thickness. Coronal sections containing the midbrain and striatum were collected in ice-cold PBS. Sections were blocked in PBS containing 0.25% TritonX-100 and 10% goat serum (PBST) for 1 h at room temperature. Slices were incubated overnight in primary antibody in PBST at 4 °C. The following primary antibodies were used: rabbit polyclonal anti-HA (1:500, A40, RRID:AB_10693385) or mouse monoclonal anti-HA (1:500, A12, RRID:AB_2565006), and guinea pig polyclonal anti-TH (1:1000, A111, RRID:AB_2619897). Slices were washed 3x in PBST followed by 2 h incubation with secondary antibody in PBST for 2 h at room temperature in the dark. The following secondary antibodies were used: goat anti-rabbit IgG Alexa 488 (1:500, S5, RRID:AB_2576217), goat anti-mouse IgG1 Alexa 488 (1:500, S7, RRID:AB_2535764), goat anti-guinea pig IgG Alexa 633 (1:500, S34, RRID:AB_2535757). Slices were washed again 3x with PBST before being mounted on glass slides with Fluoromount-G (Southern Biotech 0100-01). Stained slices were then imaged on an Olympus FV1000 confocal microscope with a 60x objective. Images were pseudo-colored in ImageJ for display. All image acquisition was done in comparison to an uninfected control imaged at the same time to assess background fluorescence. Representative images were brightness and contrast adjusted to facilitate inspection, and these adjustments were made identically for images within the same experiment. All images in Fig. 1C except for those of mice expressing ELKS2β-BirA were taken at the same time. Images of mice expressing ELKS2β-BirA were taken in a separate session and compared to their own uninfected control to confirm signal specificity. Three to five images per virus condition were taken during each imaging session, and the experiment was repeated in at least three mice per condition.

### Biotin affinity purification

Biotin affinity purifications were adapted from previously established methods (Uezu et al., 2016). Between two (for pilot experiments) and 12 mice (for mass spectrometry analyses) aged 65 to 100 days, previously injected with AAVs for BirA fusion protein expression and subjected to subcutaneous biotin injections were deeply anesthetized with isoflurane and decapitated. Brains were collected in ice cold PBS and striata were dissected out. Except for those used in pilot experiments, striata were then flash frozen and stored at -80 °C until further processing. For pilot experiments, two to four dissected striata were immediately homogenized. For mass spectrometry, five to six dissected striata were homogenized at a time. Homogenization was performed using a glass-Teflon homogenizer in 1 ml of homogenizing buffer (50 mM HEPES, 150 mM NaCl, 1 mM EDTA + mammalian protease inhibitor cocktail (Sigma CAT# P8340)) with 30 slow strokes on ice. An appropriate volume of 5x lysis buffer (1% SDS, 5% Triton X-100, 5% deoxycholate in homogenizing buffer) was added to the homogenized tissue (working concentration 0.2% SDS, 1% Triton X-100, 1% deoxycholate) and incubated while rotating at 4 °C for 1 h. Because the mass spectrometry conditions were split into two batches of five to six striata each for homogenization, the batches were combined after lysis into a single tube. Lysed samples were then sonicated twice for 10 s each at 4 °C with a Branson Sonifier 450. Sonicated samples were centrifuged at 15,000 x g for 15 min at 4 °C on a tabletop centrifuge. The cleared supernatant was removed and added to open-top polycarbonate tubes (Beckman CAT# 343778) and centrifuged in a tabletop ultracentrifuge (Beckman Rotor TLA120.2) for 1 h at 100,000 x g. After ultracentrifugation, SDS from 0.4% and/or 5% SDS stock solutions was added to the cleared supernatant to make a final SDS concentration of 1% and final volume of 1.5 ml. The sample was boiled at 95 °C for 5 min and allowed to cool to room temperature. Neutravidin agarose beads (Thermo CAT# 29200) were washed three times in binding buffer (1% SDS, 1% Triton X-100, 1% deoxycholate in homogenizing buffer). 20 µl washed neutravidin beads were added to each cooled sample and incubated for 16 h at 4 °C. After neutravidin bead incubation, the flowthrough was removed and the beads were transferred to Protein Lo-bind tubes (Eppendorf CAT# 022431081). Beads were washed twice with 500 µl 2% SDS in H2O, twice with 500 µl 1% Triton X-100, 1% deoxycholate, 25 mM LiCl in H2O, and twice in 500 µl 1 M NaCl. For each washing step, beads were pelleted by spinning 500 x g for 2 min in tabletop centrifuge at 4 °C in between each wash. Bead pellets were then washed five times in 500 µl 50 mM ammonium bicarbonate in water with spinning 500 x g for 2 min between steps. After the final wash the bead pellet was stored at -20 °C until next steps (either Western blot or PC removal).

### Western blot after biotin pulldown

Neutravidin bead pellets were incubated in 60 µl of 1x SDS-PAGE loading buffer and boiled for 10 min at 95 °C. The sample was spun at 13,000 x g on a tabletop centrifuge for 1 min to pellet the beads. 15 µl of the supernatant were loaded on an SDS-PAGE gel. 1% of the total input used for the binding reaction (cleared lysate just before addition of the neutravidin beads) was also loaded onto the gel and transferred on a nitrocellulose membrane for comparison. Membranes were blocked in 10% milk, 5% goat serum in Tris-buffered saline + 0.1% Tween-20 (TBST) for 1 h at room temperature and then incubated overnight at 4 °C in primary antibody diluted in antibody binding solution (blocking solution diluted 1:1 with TBST). Primary antibodies used: mouse monoclonal anti-HA (1:500, LAC A12, RRID:AB_2565006), rabbit polyclonal anti-red fluorescent protein (RFP) (1: 500, LAC A81, RRID:AB_2209751), rabbit polyclonal anti-pyruvate carboxylase (1:500, LAC A252, RRID:AB_11016707), mouse monoclonal anti-β-actin (1:10,000, LAC A127, RRID:AB_476692). Membranes were washed three times with TBST and incubated in HRP-conjugated secondary antibodies for 1 h at room temperature. Membranes were washed three times in TBST and enhanced chemiluminescence followed by exposure to film was used to visualize protein bands.

### Pyruvate carboxylase depletion

For samples being submitted to mass spectrometry, frozen neutravidin bead pellets were thawed on ice. Proteins were eluted by incubation in 500 µl RapiGest elution buffer (0.1% RapiGest, Waters CAT# 1866001861, in 2 mM biotin, 50 mM ammonium bicarbonate) in water for 2 h at 60 °C, while shaking. Neutravidin beads were pelleted by spinning at 18,000 x g for 5 min at 4 °C and the supernatant was moved to a new tube. To prepare anti-pyruvate carboxylase (PC) antibody-conjugated beads, Protein G Sepharose beads (GE Healthcare CAT#17-0618-01) were washed 3x in RapiGest elution buffer and conjugated to anti-PC antibodies (Novus, A252, RRID:AB_11016707) by incubating 3 μl of antibody per 20 µl of beads for 1 h at 4 °C on a rotator. Conjugated beads were spun down at 1,000 x g for 2 min, the supernatant was removed and beads were diluted 1:1 with fresh RapiGest elution buffer to make a 50% slurry. 20 µl sepharose beads (40 µl of a 50% slurry) were added to the 500 µl of eluted protein solution of the biotin affinity purification and incubated for 1 h at 4 °C on a rotator. PC-conjugated beads were removed by spinning the sample at 18,000 x g for 5 min and removing the supernatant. The proteins in the PC-depleted supernatant were then precipitated using trichloroacetic acid (TCA) precipitation. One volume of 100% TCA was added to four volumes of the PC-depleted supernatant and inverted several times before incubation on ice for 10 min. Tubes were then spun at 20,000 x g for 10 min at 4 °C. The supernatant was removed leaving behind a white protein pellet. The pellet was air dried and stored at -20 °C until analysis by mass spectrometry. In pilot experiments, we assessed the efficiency of PC depletion. Before PC depletion 28% and 10% of the detected peptides were from PC and PCCA, respectively. After depletion, 3% and 1.7% were from PC and PCCA, establishing efficient depletion and significantly enhancing the sensitivity of the mass spectrometry for all other proteins.

### Mass spectrometry

Liquid chromatography tandem mass spectrometry (LC-MS/MS) was performed by the Taplin Mass Spectrometry Facility at Harvard Medical School. Either the neutravidin bead pellet (some pilot experiments) or the TCA-precipitated protein pellet after PC depletion (all experiments to assess release site composition) were submitted to the Taplin facility. The pellet was subjected to 5 ng/µl trypsin digest overnight at 37 °C and dried until further analysis. On the day of analysis, the samples were reconstituted in 2.5% acetonitrile, 0.1% formic acid and loaded via a Famos auto sampler (LC Packings, San Francisco CA) onto the column after column equilibration. Peptides were eluted with increasing concentrations of 97.5% acetonitrile, 0.1% formic acid. As peptides were eluted, they were subjected to electrospray ionization and added to a LTQ Orbitrap Velos Pro ion-trap mass spectrometer (Thermo Fisher Scientific). Peptides were detected, isolated, and fragmented to produce a tandem mass spectrum of specific fragment ions for each peptide. Peptide sequences (and hence protein identity) were determined by matching protein databases with the acquired fragmentation pattern by the SEQUEST software program (Thermo Fisher Scientific). The data were filtered to establish a false discovery rate between one and two percent by using a database of mouse protein sequences made up of half real protein sequences and half reversed sequences. Independent of whether the results were run against a mouse or a rat database, the number of peptides identified for bait proteins (which were made from rat cDNA) were similar.

### Analsyes of mass spectrometry data

The control dataset is made up of four repeats that stem from two separate mass spectrometry runs (repeats 1 and 2 were run first, repeats 3 and 4 were run ∼ 1 yr later). Each mutant dataset (RIM cKO^DA^ and Syt-1 cKO^DA^) is made up of two repeats that were run in the same mass spectrometry run together with repeats 3 and 4 of the control dataset. Mitochondrial proteins were removed from the analyses using MitoCarta3.0 (Broad Institute), except for Figs. 3B and 6. Endogenously biotinylated PC and PCCA were also removed from the analyses along with neutravidin and IgGs. Peptide counts for each protein in each BirA condition (RIM^PPCP^-BirA, ELKS2β-BirA, CaVβ4-BirA, BirA-tdTomato) were first averaged for the two repeats that were acquired in the same mass spectrometry run. If the peptide count averaged from the two repeats was zero, it was assigned a value of 0.5. The average peptide count for a given protein for each of the three BirA baits was divided by the average peptide count for the same protein in the BirA-tdTomato condition. This is referred to as the fold change (FC) value. For the control dataset, the average fold change values of each mass spectrometry run were then averaged together to generate a final fold change value for every protein shown in Fig. 2; in the mutant datasets this final averaging was not performed as all data stem from a single mass spectrometry run. Proteins with at least two-fold enrichment over BirA-tdTomato in any release site-BirA condition were considered “hits”. The Venn diagrams were constructed using Cytoscape (v3.8.2) to map the circle size to fold change value. Circle size reflects average enrichment across repeats of an individual bait. For proteins enriched in multiple bait conditions, circle size corresponds to the condition with the largest average enrichment value. The log_2_ of the fold change was calculated for any protein that had at least a single peptide found for any of the BirA fusion proteins. To sort proteins for synaptic localization, either the enriched proteins (Figs. 2B, 5, S3 and S4) or all proteins (Fig. 4) were ran through the SynGO database (https://www.syngoportal.org) (Koopmans et al., 2019) and selected by their cellular component annotation. For Fig. 4, the self-biotinylated protein was removed from its own release site-BirA condition (for example, RIM was removed from analysis in the RIM^PPCP^ dataset but not in the CaVβ4 or ELKS2β-BirA datasets). For Fig. 5, the enriched proteins that possessed SynGO annotations were run through the STRING database (v11) (https://string-db.org) (Szklarczyk et al., 2019) to generate a STRING network diagram of enriched synaptic proteins.

### Synaptosome preparation and staining

Synaptosome experiments were performed according to previously established protocols (Banerjee et al., 2022; Liu et al., 2018). Mice (30 to 60 days old) were deeply anesthetized with isoflurane. Mice were decapitated and the brain was collected in ice cold PBS, and the striatum was dissected out. The striatal sections were homogenized in 1 ml of synaptosome homogenizing buffer (320 mM sucrose, 4 mM HEPES, 1x Sigma protease inhibitor cocktail (CAT#P8340), pH 7.4, with 12 slow strokes using a glass-Teflon homogenizer. One ml of homogenizing buffer was then added to the homogenate. The homogenate was spun at 1,000 x g for 10 min at 4 °C, the supernatant was pipetted into a new tube, and spun again at 12,500 x g for 15 min at 4 °C. The pellet (“P2 fraction”) was resuspended in 1 ml homogenizing buffer and homogenized again with 6 slow strokes. An additional 1 ml of homogenizing buffer was added, and the sample (∼1.5 ml) was loaded onto a sucrose gradient made up of 5 ml 1.2 M sucrose on the bottom and 5 ml of 0.8 M sucrose at the top in thin wall ultracentrifugation tubes (Beckman Coulter, Cat # 344059). The loaded gradient was ultracentrifuged at 69,150 x g for 1 h at 4 °C (SW 41 Ti Swinging-Bucket Rotor, Beckman Coulter, Cat. #331362) and the synaptosome fraction was harvested from the interface between the two sucrose layers. The synaptosome fraction was diluted 20-to 40-fold in homogenizing buffer and 1 ml was added to a Poly-D-Lysine-coated coverslip (neuVitro CAT# GG-18-1.5) in a 12-well plate and spun for 15 min at 4,000 x g. The buffer was removed and the synaptosomes adhering to the coverslips were fixed with 4% PFA in PBS for 10 min. The PFA was removed, and a solution with 3% bovine serum albumin and 0.1% Triton X-100 in PBS (TBP) was used for blocking and permeabilization for 1 h at room temperature. The following primary antibodies were used (diluted in TBP) overnight at 4 °C: goat polyclonal anti-NPTN (1:500, A253, RRID:AB_2155920), mouse monoclonal IgG2a anti-Bassoon (1:1000, A85, RRID:AB_11181058), and guinea pig polyclonal anti-TH (1:1000, A111, RRID:AB_2619897). After primary antibody incubation, the coverslips were washed three times in TBP and then incubated in secondary antibodies for 2 h at room temperature in the dark. Secondary antibodies used were: donkey anti-goat Alexa 488 (1:500, S6, RRID:AB_2534102, donkey anti-mouse Alexa 555 (1:500, S48, RRID:AB_2534013), donkey anti-guinea pig Alexa 647 (1:500, S59, RRID:AB_2340476). The stained synaptosomes were washed three more times in TBP before mounting on glass slides with Fluoromount-G (Southern Biotech 0100-01).

### Confocal microscopy and image analyses of stained synaptosomes

Fixed synaptosomes on coverslips were imaged with an oil immersion 60x objective and 1.5x optical zoom on an Olympus FV1000 confocal microscope. Raw confocal images were analyzed in a custom MATLAB program as described before (Liu et al., 2018) and the code was deposited on github (https://github.com/kaeserlabl/3DSIM_analysis _CL) (Liu, 2020). 1,000 to 2,000 synaptosomes per image were detected using Otsu intensity thresholds and size thresholds (0.2 to 1 µm^2^ for TH and 0.15 to 2 µm^2^ for NPTN and Bassoon). These threshold settings were identical for every image analyzed and used to detect Bassoon-positive (Bassoon+) ROIs, TH-positive (TH+) ROIs, and double-positive ROIs (Bassoon+/TH+). TH+ ROIs which had a Bassoon signal less than 1x the average intensity of all pixels in the image were designated as Bassoon-negative (Bassoon-). NPTN intensities within Bassoon+ TH+ or Bassoon-TH+ ROIs were quantified and a frequency histogram was plotted. The experimenter was blind to the condition during image acquisition and analyses.

### Statistics

Statistics were performed in GraphPad Prism 9. Data are displayed as individual data points, mean ± SEM, and/or frequency histograms. Significance is presented as * p < 0.05, ** p < 0.01, and *** p < 0.001. Sample sizes were determined based on previous studies, no statistical methods were used to predetermine sample size, no outliers were excluded. An unpaired two-tailed Student’s t-tests was used in Fig. S2B, a Kolmogorov-Smirnov test was used in Fig S2C, and two-way ANOVA tests followed by Bonferroni post-hoc tests were used in Figs. 4A-4C. In all figures, sample size and the specific tests used are stated in the figure legends.

---

**Figure S1.**
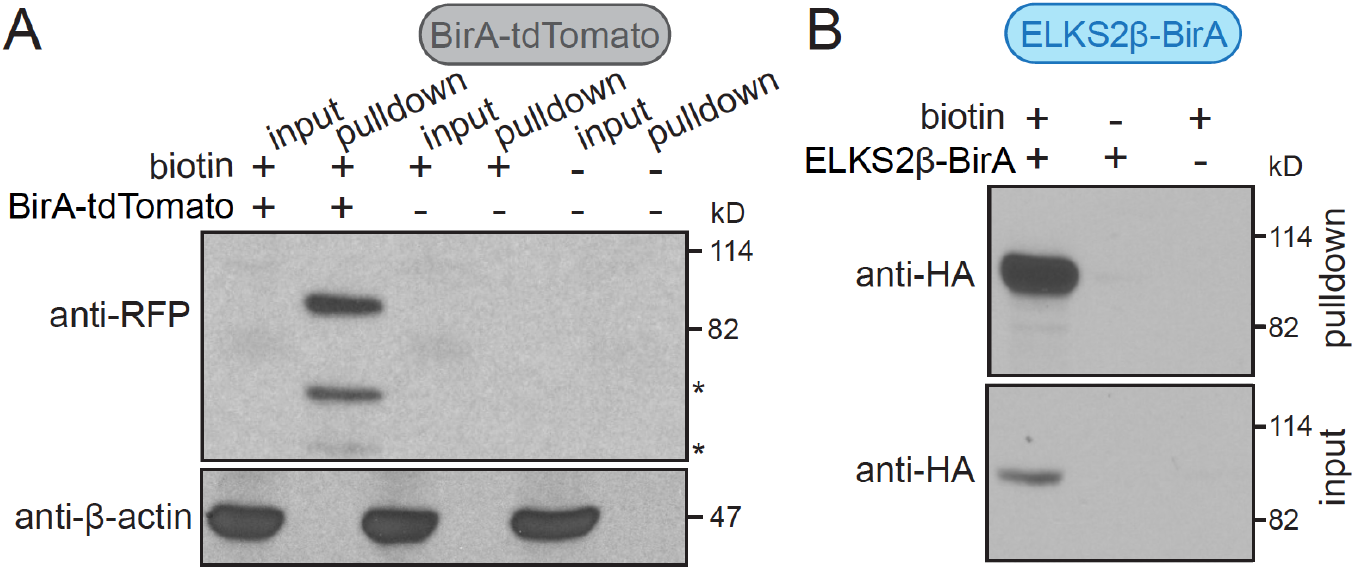
BirA fusion proteins are self-biotinylated and purified with iBioID. **(A)** Western blots of striatal lysates (input) and the pellet of the affinity-purified fractions (pulldown) are shown. The striata were harvested from mice that were injected with AAVs expressing BirA-tdTomato, or from uninjected control mice, with or without subcutaneous injection of biotin. For a timeline of the experiment, see Fig. 1A. BirA-tdTomato was detected with anti-red fluorescent protein (RFP) antibodies. The RFP bands, reflecting self-biotinylated BirA-tdTomato, were only detected in mice subcutaneously injected with biotin, but not when biotin was not supplied or when BirA-tdTomato was not expressed. β-actin was present in the brain input, but was depleted during the affinity purification. BirA-tdTomato expression is too low to be detected in the input given that dopamine axons account at most for few % of the striatal volume. The full-length BirA-tdTomato protein has a molecular weight of 91 kD. Because the smaller molecular weight bands detected with the anti-RFP antibodies (marked with *) are only present when BirA-tdTomato is enriched, they are likely degradation products. Western blots shown are from a single qualitative pilot experiment, 2 striata/condition. **(B)** Same as A, but in mice expressing ELKS2β-BirA. ELKS2β-BirA is detected with anti-HA antibodies; Western blots from a single qualitative pilot experiment, 4 striata/condition. These experiments reveal that BirA fusion proteins are self-biotinylated in vivo after biotin injections and are purified with iBioID, and they are specific enough to avoid contamination from strongly expressed, non-biotinylated proteins present in all cells, for example β-actin.

**Figure S2.**
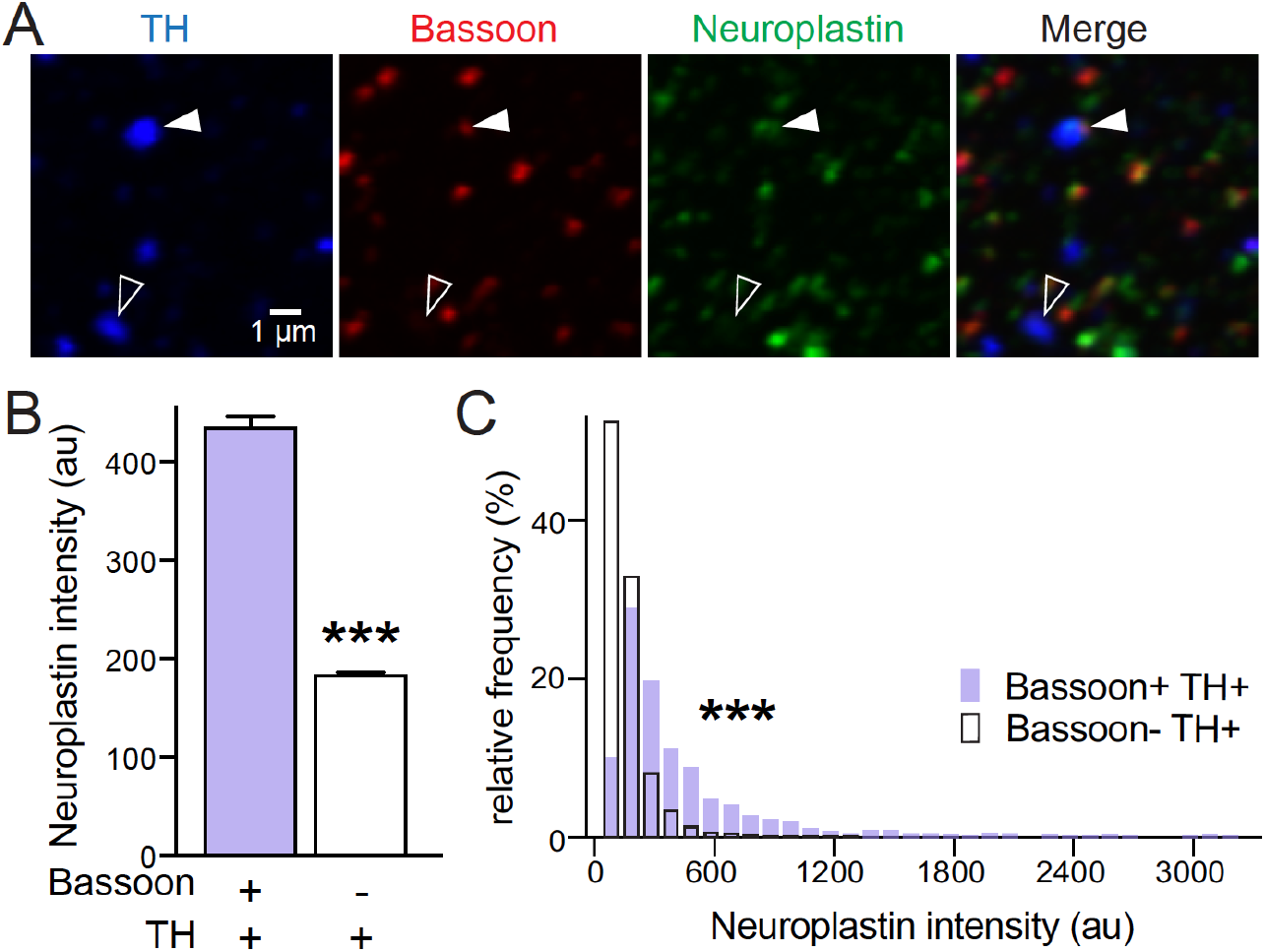
Neuroplastin is enriched in Bassoon-containing dopaminergic synaptosomes. **(A)** Representative confocal images of striatal synaptosomes stained with anti-Bassoon antibodies to mark release sites, anti-TH antibodies to label dopamine synaptosomes, and anti-Neuroplastin antibodies. Only a fraction of TH-positive synaptosomes contains the release site marker Bassoon (filled arrowhead), while many do not (outlined arrowhead) (Liu et al., 2018). **(B, C)** Quantification of the experiment shown in A. Neuroplastin intensity within TH-positive (TH+) and TH-negative (TH-) synaptosomes was quantified (B), and an intensity histogram was constructed (C); Bassoon+ TH+, 1316 synaptosomes/30 images/3 mice; Bassoon-TH+, 5434/30/3. Data are shown as mean ± SEM. *** p <.0001 as assessed by an unpaired t-test (B) or a Kolmogorov-Smirnov test (C).

**Figure S3.**
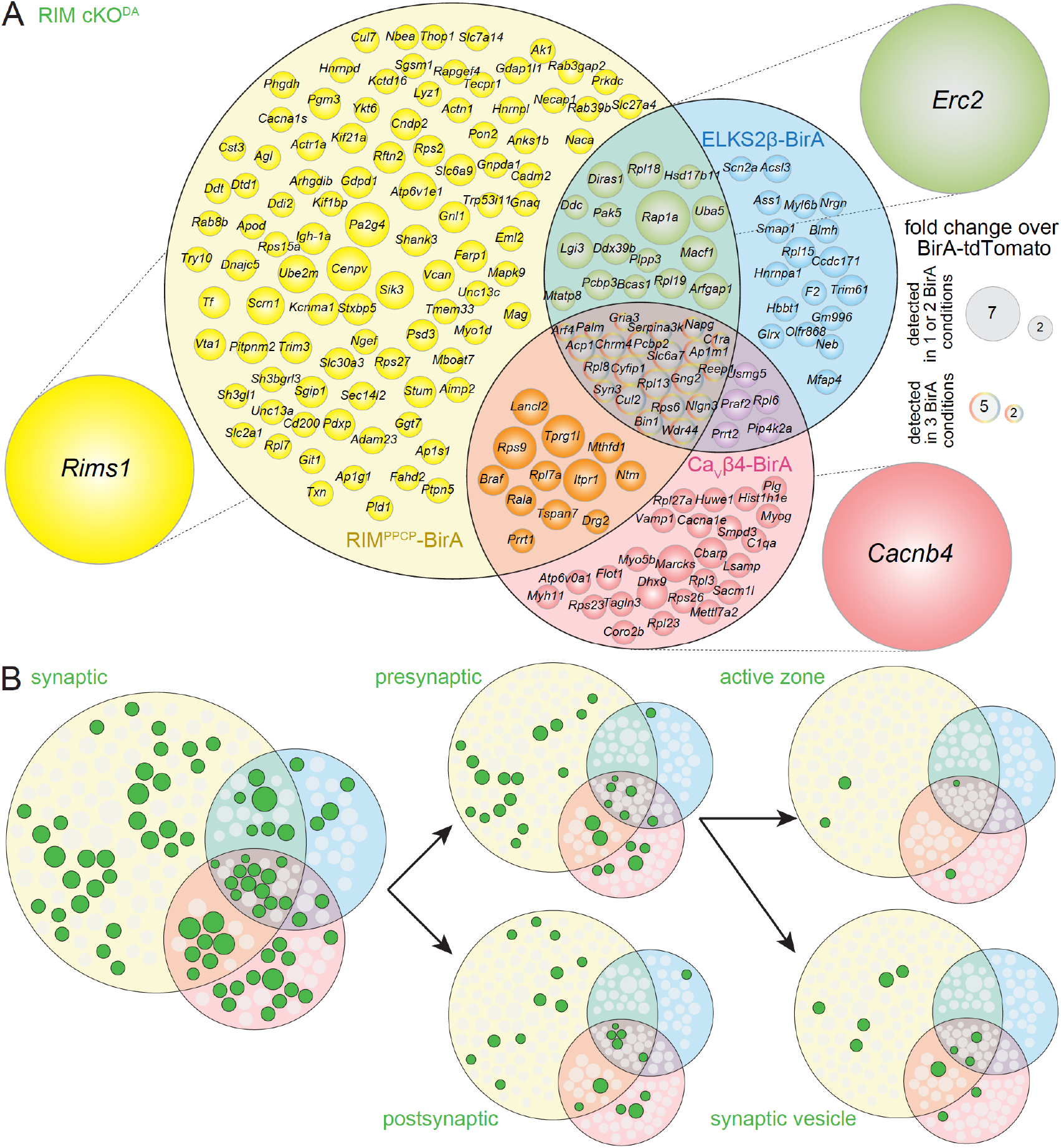
Venn diagrams for the RIM cKO^DA^ dataset. **(A)** Venn diagram listing genes that encode proteins enriched ≥ 2-fold over BirA-tdTomato with RIM^PPCP^-BirA (yellow), ELKS2β-BirA (blue) or CaVβ4-BirA (red) iBioID baits in RIM cKO^DA^ mice. Proteins used as BirA baits are shown outside the Venn diagram in the color of the corresponding part of the diagram. Enrichment is shown as the average of two independent repeats run in a single mass spectrometry session. Circle size reflects average enrichment across repeats (key on the right). For proteins enriched in multiple bait conditions, circle size corresponds to the bait condition with the largest enrichment. **(B)** Enriched proteins from (A) that have cellular compartment annotations in the SynGO database (Koopmans et al., 2019) are colored in green and classified into increasingly specific SynGO sub-categories, sub-categorizations are not mutually exclusive.

**Figure S4.**
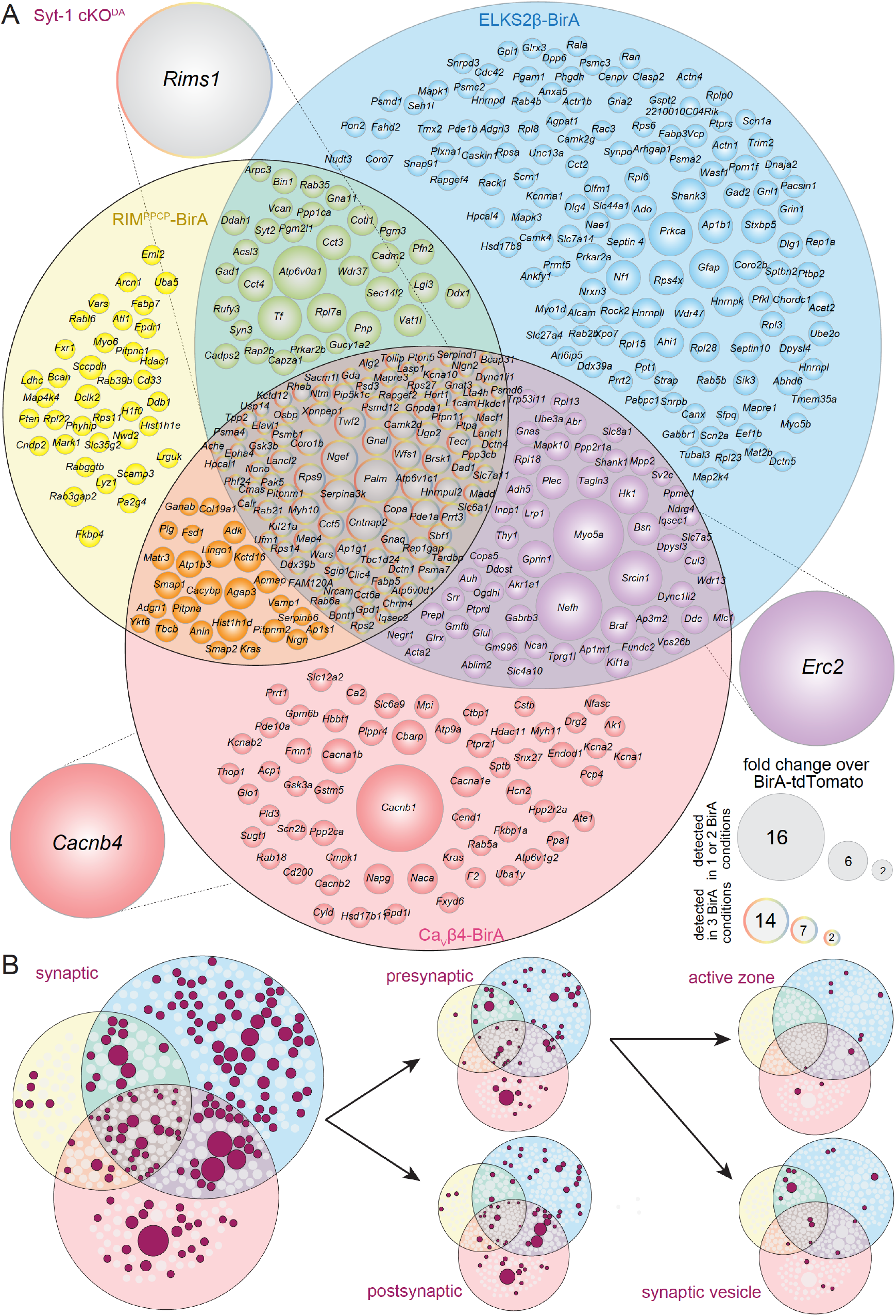
Venn diagrams for the Syt-1 cKO^DA^ dataset. **(A)** Venn diagram listing genes that encode proteins enriched ≥ 2-fold over BirA-tdTomato with RIM^PPCP^-BirA (yellow), ELKS2β-BirA (blue) or CaVβ4-BirA (red) iBioID baits in Syt-1 cKO^DA^ mice. Proteins used as BirA baits are shown outside the Venn diagram in the color of the corresponding part of the diagram. Enrichment is shown as the average of two independent repeats run in a single mass spectrometry session. Circle size reflects average enrichment across repeats (key on the right). For proteins enriched in multiple bait conditions, circle size corresponds to the bait condition with the largest enrichment. **(B)** Enriched proteins from (A) that have cellular compartment annotations in the SynGO database (Koopmans et al., 2019) are colored in maroon and classified into increasingly specific SynGO sub-categories, sub-categorizations are not mutually exclusive.

## Notes

### Competing Interest Statement

The authors have declared no competing interest.

